# Protein Kinase C positively regulates peroxisome biogenesis by promoting peroxisome-endoplasmic reticulum interaction

**DOI:** 10.1101/2025.01.21.634043

**Authors:** Anya Borisyuk, Charlotte Howman, Sundararaghavan Pattabiraman, Daniel Kaganovich, Triana Amen

**Affiliations:** Global Health Institute, Faculty of Life Sciences, Ecole Polytechnique Fédérale de Lausanne, Lausanne, Switzerland; School of Biological Sciences, University of Southampton, Southampton, UK; 1BasePharma, Israel; 2Prime Ltd, UK

**Keywords:** Peroxisome, Protein Kinase C, GSK3β, PEX11b, VAPB, contact sites, peroxisome biogenesis, peroxisome division, signaling

## Abstract

Peroxisomes carry out a diverse set of metabolic functions, including oxidation of very long-chained fatty acids, degradation of D-amino acids and hydrogen peroxide, and bile acid production. Many of these functions are upregulated on demand, therefore cells control peroxisome abundance, and by extension peroxisome function, in response to environmental and developmental cues. The mechanisms upregulating peroxisomes in mammalian cells have remained unclear. Here we identify a signaling regulatory network and a mechanism that coordinate cellular demand for peroxisomes and peroxisome abundance by regulating peroxisome proliferation. We show that protein kinase C (PKC) promotes peroxisome PEX11b-dependent formation. PKC activation leads to an increase in peroxisome formation, promoting peroxisome-ER contact site formation through inactivation of GSK3β. We show that removal of VAPA and VAPB impairs peroxisome biogenesis and PKC regulation. Inhibition of PKC reduces peroxisome-ER interactions, leading to a decrease in peroxisome abundance. During neuronal differentiation, active PKC leads to a significant increase in peroxisome formation. We propose that peroxisomal regulation by transient active PKC signaling enables rapid and fine-tuned responses to the need for peroxisomal activity.

## Introduction

The peroxisome is a small organelle with a large functional repertoire ^1–3^. Prominent examples of peroxisomal function are alpha- and beta-fatty acid oxidation, bile acid and ether-lipid synthesis, degradation of D-amino acids and leukotrienes, glyoxylate cycle and methanol metabolism in yeast, bioluminescence in fireflies, plant hormone synthesis, and degradation of hydrogen peroxide ^2–15^. In addition to their numerous metabolic functions, peroxisomes are increasingly seen as signaling regulators ^16–20^. Several kinases, (e.g. the NDR2 kinase regulator of ciliogenesis) contain peroxisome localization signals ^21, 22^. The peroxisomal membrane has been shown to recruit multiple signaling regulators, including the tuberous sclerosis complex (TSC1/2) regulator of the target of rapamycin (mTOR) kinase pathway, which binds to peroxisomal proteins PEX19 and PEX5 ^16–20^.

Due to the diversity of peroxisomal functions, the cellular demand for peroxisomes depends on specific conditions, such that peroxisomes must be rapidly mass-produced during specific instances of stress adaptation and development ^7, 23, 24^, necessitating precise regulation. Disruption of peroxisomal biogenesis adversely affects downstream cellular adaptive functions, including autophagy, cholesterol metabolism, immune response, and ciliogenesis ^20, 22, 25–29^. Not surprisingly, therefore, multiple human disorders result from defects in peroxisome biogenesis ^2, 30–32^. Due to the transient “on demand” need for peroxisomal metabolism, the key to understanding the role of peroxisomes in cell biology and disease is to investigate the mechanisms regulating peroxisomal biogenesis.

Peroxisomes form through two main pathways: fission of pre-existing peroxisomes ^33–38^, and *de novo* (from intracellular endoplasmic reticulum- and mitochondria-derived membranes) ^35, 36, 39–44^. The peroxisome biogenesis machinery consists of peroxins (PEX), which are conserved in fungi, plants, and animals ^34, 45–48^. Deletion of PEX3, PEX19 and PEX16 results in conditionally viable cells lacking peroxisomes ^34, 39, 41, 43, 44, 47, 49–52^. Peroxisome division, on the other hand, is initiated by the PEX11 protein family, which orchestrate a multistep process that involves peroxisome membrane elongation, and prime assembly of additional peroxisome division components, including MFF, FIS1, and DNM1L 53-57. Both peroxisome biogenesis mechanisms are initiated by several non-mutually exclusive pathways: transcriptional peroxisome proliferator receptor alpha (PPARα) increases the level of division factors, peroxisome tethering to the endoplasmic reticulum (ER) through a recently defined membrane contact site that is formed between acyl-coenzyme A-binding domain protein 5 (ACBD5) and the ER protein vesicle-associated membrane protein-associated protein (VAPB)^58, 59^ potentially sources the membrane, and an increase in peroxisome function in the presence of peroxisomal substrates promotes peroxisome formation via unknown mechanissms. The latter phenomenon of increased peroxisome number in response to increased peroxisomal function is known as “metabolic control of peroxisome abundance (MCPA)” ^23, 60–63^. Recently, Schrader/Costello groups demonstrated how peroxisome-endoplasmic reticulum membrane contacts are negatively regulated by glycogen synthase kinase beta (GSK3β) that phosphorylates ACBD5 preventing interaction with VAPB and reducing contacts^58, 64, 65^. We showed that abolishing the peroxisome-ER contacts through knockout of VAPB and its homologue VAPA leads to a significant defect in peroxisome biogenesis^66^.

Here, using a focused tool compound kinase inhibitor screen, we identify positive and negative regulators of peroxisome abundance, including PKC as a major positive driver. We show that active PKC inhibits GSK3β, promoting peroxisome-ER tethering and peroxisome biogenesis, promoting ACBD5-VAPB interaction^58, 67, 68^. We further show that PKC regulation is dependeend on PEX11b and VAPs. Inhibition of PKC reduces peroxisome-ER contacts, peroxisome abundance, and function. Our study shows how PKC can rapidly regulate peroxisome-ER interaction, peroxisome formation, and peroxisome function.

## Results

### Tool Compound Kinase Inhibitor screen reveals signaling regulators of peroxisome abundance

We postulated that a sensitive and tractable assay of peroxisome abundance was the key to discovering regulators of peroxisome formation and, by extension, peroxisome function. We therefore developed a new tool for quantifying peroxisomes in live cells. We used CRISPR/Cas9 gene editing to genomically fuse GFP to the C-terminus of the transmembrane peroxisomal protein PMP70 in HEK293T cells (Figure 1A; Figure S1A and S1B). PMP70-GFP localized to peroxisomes, peroxisomal protein import was not impaired, and peroxisomes in the endogenously tagged cells were comparable to the wildtype cells (Figure S1C, S1D, and S1E). Considering the prominent role of kinase signaling in metabolic regulation, we set out to identify kinase modulators of peroxisome abundance. We opted for a small molecule “tool compound” library of 152 kinase inhibitors, which we viewed as preferable to gene manipulation-based screening given the pleiotropic and essential nature of regulatory kinase function (Figure 1B; Figure S1F, Figure S2, Figure S3, and Supplementary Table 1). Peroxisome cellular density (abundance) was calculated as the number of peroxisomes per square micron of the cytoplasmic space in confocal microscopy images, using the overexpression of PEX19 peroxisome biogenesis factor as a positive control^49^, and overexpression of PEX3 as a negative control (Figure 1C) ^69, 70^. We found that 16 kinase inhibitors reduced, and 5 upregulated, the number of peroxisomes in HEK293T cells (Figure S1F and S1G). We performed a secondary screen of the hits in untagged human fibroblasts (Figure 1D), validating 10 compounds that reduced and 2 that upregulated the number of peroxisomes (Figure 1E). We then associated the molecules to the kinases they inhibit and mapped the kinases on the human kinome (Figure 1F). Positive regulators (whereby inhibition of the kinase reduces peroxisome number) included: protein kinase C (PKC), TGFβR, MEK1/2, ERK2, PDHK, IGF1R, and ALK4/7. Negative regulators (whereby inhibition of the kinase increases peroxisome numbers) were CK2 and IKKβ (Figure 1F). Two different tool compounds targeted the TGFβ and PKC pathways. Interestingly, TGFβ is a known positive regulator of peroxisome proliferation ^71^, underlining the broad coverage of our tool compound screen. We elected to further focus on the previously uncharacterized role of PKC in regulating peroxisome abundance.

**Figure 1.**
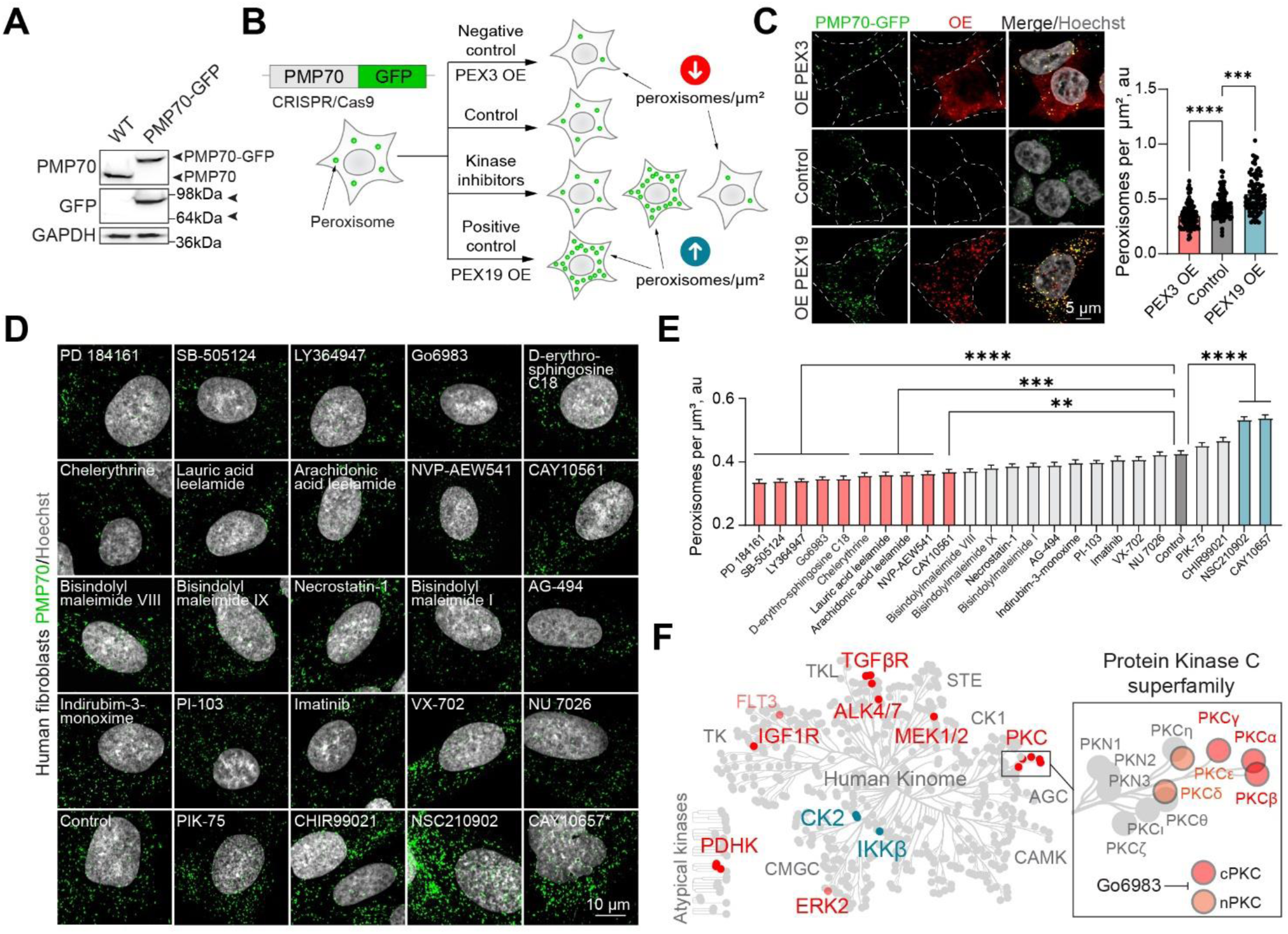
Kinase Inhibitor screen reveals signaling regulators of peroxisome abundance **(A)** Western blot of WT and CRISPR/Cas9 PMP70-GFP HEK293T cells. Arrow heads indicate the size shift of the tagged PMP70-GFP. **(B)** Schematic of the peroxisome biogenesis regulators screen. CRISPR/Cas9 PMP70-GFP HEK293T cells were incubated in control or 1μM of small molecules for 2 days, PEX3 overexpression was used as a negative control, and PEX19 overexpression was used as a positive control. Refer to Figure S1F to G, S2, and S3 for the screen details. **(C)** Confocal microscopy of CRISPR/Cas9 PMP70-GFP HEK293T cells overexpressing PEX3-myc, flag-PEX19, or an empty vector. Quantification shows the number of peroxisomes per square micron of the cytoplasm in the 2D confocal image, mean± SEM, N=100, *** - p<0.001, **** - p<0.0001. **(D-E)** Confocal microscopy of peroxisomes in human primary fibroblasts AFF11 treated with indicated kinase inhibitors for 2 days. Peroxisomes were visualized using PMP70 antibody, nuclei were stained with Hoechst (10μg/ml). Representative images are shown, Scale bar - 10μm, * - abnormal nuclear morphology. (E) Quantification shows the number of peroxisomes per square micron of the cytoplasm in the flattened 3D image (indicated as cubic micron), mean± SEM, N=100, ** - p<0.01, *** - p<0.001, **** - p<0.0001. **(F)** Identified kinase inhibitors plotted on the human kinome network ^145, 146^. Positive regulators (inhibition decreases number of peroxisomes) are indicated in red, negative regulators (inhibition increases the number of peroxisomes) are indicated in blue. Protein Kinase C superfamily is shown in the inlet, cPKC – classical PKC, nPKC – novel PKC isoforms, G06983 inhibits indicated PKC isoforms.

### Protein Kinase C delta positively regulates peroxisome abundance

First, we analyzed which PKC isoforms are inhibited by the small molecules that we identified. Go6983 PKC inhibitor prevents kinase activity as well as the steric PKC interaction with the substrates ^72^. Chelerythrine and D-erythrosphingosine C18 are potent inhibitors of PKC^73, 74^. The human PKC superfamily consists of classical (cPKC): α, βI, βII, γ; novel (nPKC): δ, ε, θ and η: atypical (aPKC): ζ and ι / λ: and PKC-related kinase (PKN) isoforms ^75, 76^. Identified molecules are not specific to one isoform, and inhibit classical (cPKC: α, β. and γ), novel (nPKC δ), and at least one atypical (aPKC : ζ), inhibition of these PKC isoforms resulted in a decrease of peroxisome abundance (Figure 1E). We, therefore decided to test whether PKC activation by overexpression of one isoform is sufficient to trigger peroxisome proliferation. First, we overexpressed a representative isoform of each class - classical PKC(α), novel PKC(δ), and atypical isoform of PKC(ζ) fused to mCherry and mCherry as a control and measured peroxisome number (Figure 2A-B). Only PKCδ overexpression was able to induce peroxisome proliferation (Figure 2A; Figure S4A). Both cPKC and nPKC bind to and are activated by phorbol esters (PMA)^77, 78^. We incubated human fibroblasts with PMA, which resulted in a significant increase in peroxisome number (Figure 2C). Together, these data argue that PKCδ is a positive regulator of peroxisome abundance in human cells. To independently test that inhibition of PKC affects peroxisome biogenesis we used a radioactive assay of peroxisomal fatty acid oxidation that we previously established^66^. As a control we generated PEX19 KO that lack peroxisomes (Figure 2D)^66^. Inhibition of PKC resulted in a significant reduction of peroxisome function, but did not inhibit it completely (Figure 2D), consistent with a reduction of peroxisome numbers. PKC, therefore, regulates peroxisome biogenesis – formation and function of peroxisomes. Next, investigated the mechanism of this new role of PKC in peroxisome biology.

**Figure 2.**
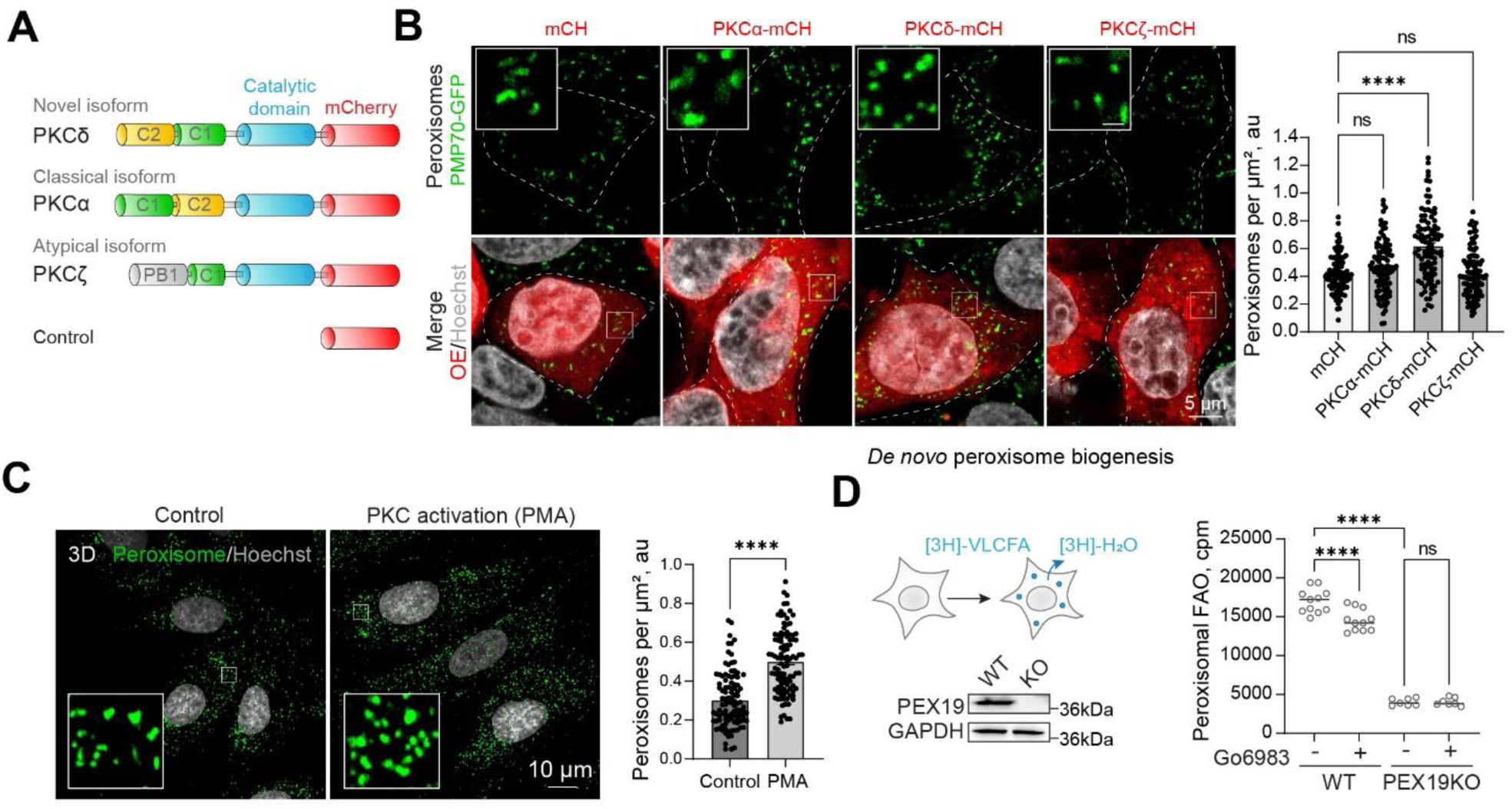
Protein Kinase C Delta positively regulates peroxisome abundance **(A)** Schematic of different PKC isoforms used in (B) **(B)** Confocal microscopy of HEK293T CRISPR/Cas9 PMP70-GFP cells overexpressing (OE) PKCα-mCherry, PKCδ-mCherry, or PKCζ-mCherry. Quantification shows number of peroxisomes per square micron of the cytoplasm in the 2D confocal image, mean± SEM, N=100, **** - p<0.0001. Scale bar - 5μm Refer to Figure S5H. **(C)** Confocal microscopy of peroxisomes in human primary fibroblasts AFF11 treated with PMA (0.5μM) for 1 day. Peroxisomes were visualized using PMP70 antibody, nuclei were stained with Hoechst (10μg/ml). Representative images are shown, Scale bar - 10μm. Quantification shows number of peroxisomes per square micron, mean± SEM, N=103, **** - p<0.0001. Refer to Figure S5I. **(D)** Radioactive peroxisomal FAO measurement using 3H-docosanoic acid in HEK293T WT or PEX19KO in control or Go6983 (5μM for 48hours) conditions. Quantification shows the number of counts per minute, mean± SEM, N=6-12, *** - p<0.001, ****

### Protein Kinase C induces PEX11b-dependent peroxisome formation

Peroxisome abundance is maintained by peroxisome biogenesis by *de novo* and division pathways, and peroxisome degradation ^35, 39, 43, 45, 53, 79–82^. To understand how PKC activation results in an increased number of peroxisomes, we first reproduced the PEX19 functional complementation assay ^41^ in HEK293T cells (Figure 3A). This assay monitors *de novo* peroxisome formation by complementing PEX19 KO cells with PEX19 and measuring peroxisome formation. We constructed CRISPR/Cas9 PEX19 KO cells, that lack peroxisomes^66^. In line with published observations, it takes more than 24 hours to restore peroxisomes *de novo* ^36, 41^ (Figure 3B). Peroxisomes appear on the second day after transfection and continue to proliferate, with a 70% restoration on day 4 (Figure 3B-D). PKC inhibition did not change the *de novo* peroxisome formation. Note, that here we measured the ratio of cells that restored peroxisomes among the cells expressing PEX19 (the density of peroxisomes in single cells was not assayed). Similarly, PKC activation with PMA did not affect *de novo* peroxisome formation, however, a long exposure to PMA is inhibitory to PKC, as active PKC is degraded ^83^. Peroxisome division is thought to be a more frequent event than peroxisome *de novo* formation ^84^, and it takes considerably less time. Peroxisome proliferation was upregulated with a short 2-hour exposure to PMA that activates PKC, pointing towards the proliferation pathway as a point of PKC regulation of peroxisome abundance (Figure 3E).

**Figure 3.**
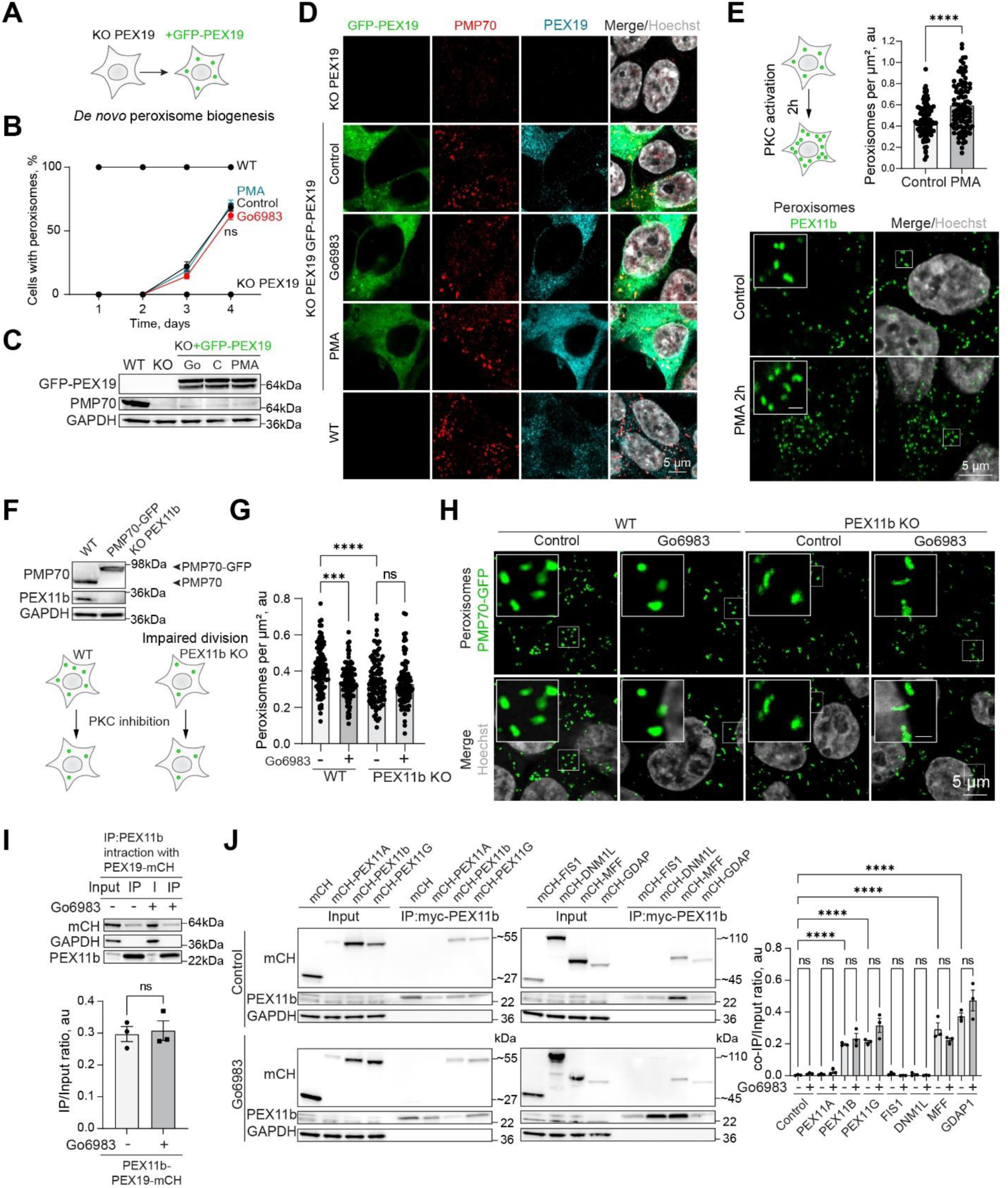
Protein kinase C regulation depends on PEX11b **(A)** Schematic of PEX19 complementation assay. **(B-D)** Confocal microscopy of the peroxisomes in HEK293T WT, PEX19 KO, and PEX19 KO cells overexpressing flag-GFP-PEX19 in control, Go6983 1μM, and PMA 0.5μM conditions. Cells were fixed and visualized on identified days. (B) Quantification shows the ratio of cells that restored peroxisomes among the transfected cells. Scale bar -5μm. (C) Western blot of PMP70 expression on day 4 of the complementation assay. **(E)** Confocal microscopy of peroxisomes in HEK293T CRISPR/Cas9 flag-GFP-PEX11b cells treated with 0.5μM PMA for 2 hours. Nuclei were stained with Hoechst (10μg/ml). Representative images are shown, Scale bar - 5μm, inlet - 1μm. Quantification shows number of peroxisomes per square micron of the cytoplasm in the 2D confocal image, mean± SEM, N=100, **** - p<0.0001. **(F)** Western blot of PEX11b and PMP70 in the WT and CRISPR/Cas9 PMP70-GFP PEX11b KO HEK293T cells. Arrow heads indicate the size shift of the tagged PMP70-GFP. **(G-H)** Confocal microscopy of peroxisomes in HEK293T CRISPR/Cas9 PMP70-GFP WT and PEX11b KO cells grown in control or Go6983 5μM conditions. Nuclei were stained with Hoechst (10μg/ml). Representative images are shown, Scale bar - 5μm, inlet - 1 μm. Quantification shows the number of peroxisomes per square micron of the cytoplasm in the 2D confocal image, mean± SEM, N=100, *** - p<0.001, **** - p<0.0001. **(I)** Co-Immunoprecipitation of mycPEX11b in control and PKC inhibition conditions (Go6983 5μM, 24 hours). Western blot of mycPEX11b and mCherry-PEX19 in the WT HEK293T cells. Quantification shows the IP to Input (I) ratio, mean± SEM, N=3. **(J)** Co-Immunoprecipitation of mycPEX11b and peroxisome division factors in control and PKC inhibition conditions (Go6983 5μM, 24 hours). Western blot of mycPEX11b and mCherry-, mCherry-PEX11A, PEX11b, PEX11G, FIS1, DNM1L, MFF, or GDAP1 in the WT HEK293T cells. Quantification shows the IP to Input (I) ratio, mean± SEM, N=3, **** - p<0.0001.

Peroxisome proliferation is controlled by PEX11b - a key peroxisomal proliferation factor, responsible for early membrane remodeling, elongation, and association with the rest of the division machinery that is shared with mitochondria ^38, 53, 54^. In order to understand whether PKC regulates PEX11b-induced peroxisome division we constructed a CRISPR/Cas9 PEX11b KO (Figure 3F). PEX11b KO has a reduction in peroxisome abundance as was previously shown, and it did not respond to PKC inhibition by further reduction of peroxisome numbers (Figure 3G and 3H) ^54, 85^.

Finally, to rule out PKC-driven regulation of the PPARα peroxisomal transcriptional regulator ^86^, we measured PPARα levels following PKC inhibition and activation. PKC inhibition had no effect on PPARα, whereas PKC activation with PMA showed a modest increase in PPARα levels, though not PPARα activity (Figure S4B-E)^87, 88^. To also rule out a role for pexophagy ^89^, we silenced the NBR1 receptor, which did not affect peroxisome abundance over a two day period (Figure S4F, S4G), confirming that PKC does not upregulate peroxisomes by inhibiting their degradation. Collectively these data points to the peroxisome division pathways as a route for PKC regulation of peroxisome abundance. Next, we investigated the mechanism of PKC regulation of peroxisome proliferation.

### PKC phosphorylation of PEX11b on S53 is not sufficient to induce peroxisome proliferation

First, to explore whether PEX11b machinery is reorganized during PKC inhibition, we tested whether PEX11b changes its association with other division factors. However, neither the interaction of PEX11b with other factors of peroxisome division machinery or PEX19 (Figure 3I-J) were significantly affected.

PEX11 phosphorylation is known to promote peroxisome division in certain yeast species (*Saccharomyces cerevisiae* and *Pichia pastoris)*, but not in others (*H. polymorpha)* ^90–92^. Therefore, we were curious whether phosphorylation of PEX11b by PKC regulates proliferation of peroxisomes in human cells. There are 6 predicted PKC phospho-sites on PEX11b according to NetPhos3.1 prediction algorithm ^93, 94^. Mass spectrometry analysis of the PEX11b in the lysate results in low peptide coverage, not sufficient for the phospho-proteomic identification. To overcome this limitation, we constructed a CRISPR/Cas9 flag-GFP-PEX11b HEK293T cell line (Figure S5A). Flag pull-down of PEX11b significantly improved PEX11b coverage (Figure S6A), and was used for targeted phospho-peptide identification (Figure S5B, Supplementary Table 2). Among the identified peptides, Ser53 is a predicted PKC target, Ser38 is an unspecified/p38MAPK target, and Ser78 was not identified by the prediction algorithm (Figure S5B; Figure S6B-C). The ratio of phosphorylated to the overall identified peptides was 4.4% for Ser53. To confirm that PKC can directly phosphorylate Ser53 of PEX11b we used a pure peptide competition assay – a mix of all the predicted PKC-phosphorylated PEX11b peptides with a known PKC substrate peptide ^95^, incubated with or without the addition of pure active PKC and subjected to phospho-mass spectrometry (Figure S5B). 4 out of 6 predicted peptides were phosphorylated to a similar extent as the PKC substrate peptide, including the Ser53 site, and none of the peptides were phosphorylated without PKC (Figure S5B-C, S6D).

We overexpressed PEX11b, PEX11b S53A (Ser53Ala), and a phosphomimetic PEX11b S53D (Ser53Asp) mutant in PEX11b KO background and assessed peroxisome number (Figure S5C-F). PEX11b S53D overexpression, and that of the WT and S53A, had no effect on peroxisome number (after correction for PEX11b expression level) (Figure S5E). Importantly, N-terminal phosphorylation of PEX11b on S38, was also shown to have no effect on peroxisome division ^96^. We also verified that other identified sites (S78, and a triple D mutant) did not affect the number of peroxisomes either (Figure S5G-H).

### PKC regulates peroxisome-ER contact sites through GSK3β inhibition

Given that PKC phosphorylation of PEX11b failed to explain its effect on peroxisome proliferation, we posited that the regulation of the peroxisome-ER interaction can affect peroxisome proliferation. This is based on the discovery of the VAPB-ACDB5 contacts^59, 68^ and our finding that KO of VAPA and VAPB results in a significant decrease in peroxisome abundance^66^. In addition to acting as a structural anchor, the VAPB-ACBD5 contact site regulates lipid transfer between the ER and peroxisomes. Since peroxisomes do not produce their own phospholipids, this activity may be essential for division-associated membrane growth^58^. To independently confirm that peroxisome-ER contact sites are required for peroxisome biogenesis (i.e. an increase in the number of peroxisomes), we used VAPB/VAPA knockout cells ^97^. There was a significant decrease in peroxisome number in the VAP KO cells, which was restored upon complementation with VAPB (Figure 4A, Figure S4H). Peroxisome abundance in VAP KO cells was significantly reduced, and not affected by PKC inhibition (Figure 4B)^66^.

**Figure 4.**
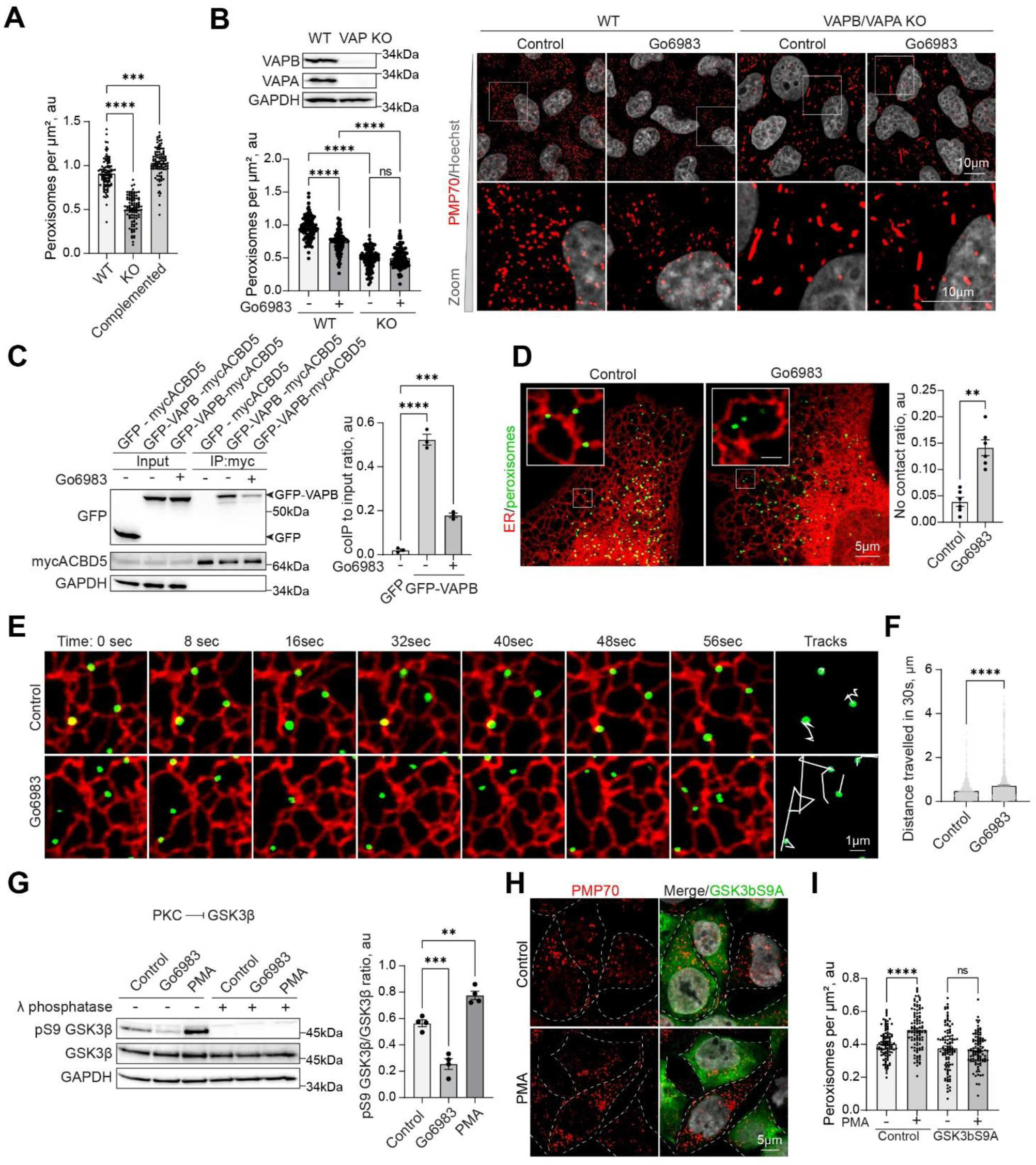
PKC regulates peroxisome-ER VAP-dependent interaction through GSK3b inhibition **(A)** Quantification of confocal microscopy of peroxisomes stained with antiPMP70 antibody in WT and VAPB/VAPA KO HeLa cells in control and GFP-VAPB overexpression conditions. Quantification shows the number of peroxisomes per micron square, mean± SEM, N=100, *** - p<0.001, **** - p<0.0001. Refer to figure S4H **(B)** Confocal microscopy of peroxisomes stained with anti-PMP70 antibody in WT and VAPB/VAPA KO HeLa cells in control Go6983 (5μM for 48hours). Quantification shows the number of peroxisomes per micron square, mean± SEM, N=100, *** - p<0.001, **** - p<0.0001. Western blot confirming the KO is shown. **(C)** Co-Immunoprecipitation of mycACBD5 in control and PKC inhibition conditions (Go6983 5μM, 24 hours), mycACBD5 and GFP or GFP-VAPB were overexpressed in HEK293T cells. Quantification shows the IP to Input ratio, mean± SEM, N=3, *** - p<0.001, **** - p<0.0001. **(D-F)** Live cell confocal microscopy of ER and peroxisomes in U2OS cells in control and Go6983 conditions. Quantification shows the ratio of peroxisomes that are not proximal or overlapping with the ER, mean± SEM, N=6 pooled from 1000 peroxisomes in at least 30 cells per condition, ** - p<0.01, and a distance travelled by a single peroxisome in 30seconds, mean± SEM, N=1000, **** - p<0.0001. **(G)** Western blot of GSK3β inhibitory S9 phosphorylation in PMA(0.5μM for 2hours) and Go6983 (5μM for 4hours) conditions. Quantification shows the ratio of phosphorylated pS9 GSK3β to non-phosphorylated GSK3β, mean± SEM, N=4, *** - p<0.001, ** - p<0.01. **(H)** Confocal microscopy of peroxisomes stained with antiPMP70 antibody in HEK293T control or overexpression of GSK3β S9A mutant in control or PMA (0.5μM for 4hours) conditions. (I) Quantification shows the number of peroxisomes per micron square, mean± SEM, N=100, **** - p<0.0001.

We then assessed peroxisome-ER interaction^58, 59^. Inhibition of PKC resulted in a decrease in the VAPB-ACBD5 interaction as evidenced by the co-immunoprecipitation of VAPB (Figure 4C), and resulted in a significant increase in a free-roaming peroxisomes, as evidenced by the lack of peroxisome-ER proximity (Figure 4D), and significant increase in mobility (Figure 4E-F; Supplementary Movies 1-6). One obvious candidate for the mechanism of this regulation was GSK3β, due to its established role in the VAPB-ACBD5 negative regulation^58, 68^. PKC negatively regulates GSK3β ^98, 99^, which, in turn, negatively regulates peroxisome contact sites with the ER ^68^. We therefore hypothesized that PKC drives peroxisome division by modulating contact site formation. We confirmed that PKC activation inhibits GSK3β, whereas inhibition of PKC leads to GSK3β activation, as assayed by measuring GSK3β S9 inhibitory phosphorylation (Figure 4G). Next, to confirm that PKC regulates peroxisome biogenesis through GSK3β inactivation, we constructed a GSK3β S9A constitutively active mutant^100^, which suppressed peroxisome proliferation in response to PMA treatment (PKC activation) (Figure 4H and 4I). Finally, analysis of our hits from the small molecule screen, revealed that inhibitor of GSK3β, CHIR99021, increased peroxisome number in HEK293T cells (Figure S1G). Taken together, these data shows that PKC can positively regulate peroxisome number through inhibition of GSK3β, by promoting peroxisome-ER interaction through known mechanisms^58, 68^, which in turn is regulating peroxisome biogenesis (Figure 4A)^66^.

### Protein kinase C regulates peroxisome abundance during neuronal differentiation

Finally, we investigated the biological relevance of PKC-regulated peroxisome division. PKC activity varies between tissues, with the highest levels detected in the brain ^101^. Further, peroxisome regulation is critical for brain development, evidenced by the neurodegenerative conditions stemming from impaired peroxisome biogenesis ^24^. We, therefore, tracked peroxisome abundance in differentiating neuronal cells. We used human neuroblastoma SH-SY5Y cells that can be differentiated into neuron-like cells that express neuronal markers after 18 days ^102^ (Figure 5A; Figure S7A and S7B). PKC activity was significantly increased in differentiating cells (Figure 5B), as was peroxisome number. We then differentiated SH-SY5Y neuronal cells in the presence of PKC inhibitor, that was added from day 10 to day 18 of neuronal differentiation. Apart from PAX6, differentiation markers, including MAP2, NeuN, and β-3 tubulin were upregulated to a similar extent in control and PKC-inhibited samples (Figure 5C; Figure S7B), but peroxisome number was decreased significantly (Figure 5D to F, Figure S7C). Similarly, we detected an increase in PKC activity and peroxisome number while differentiating iPSC to neuronal progenitor cells (Figure 5G-J; Figure S7D-F).

**Figure 5.**
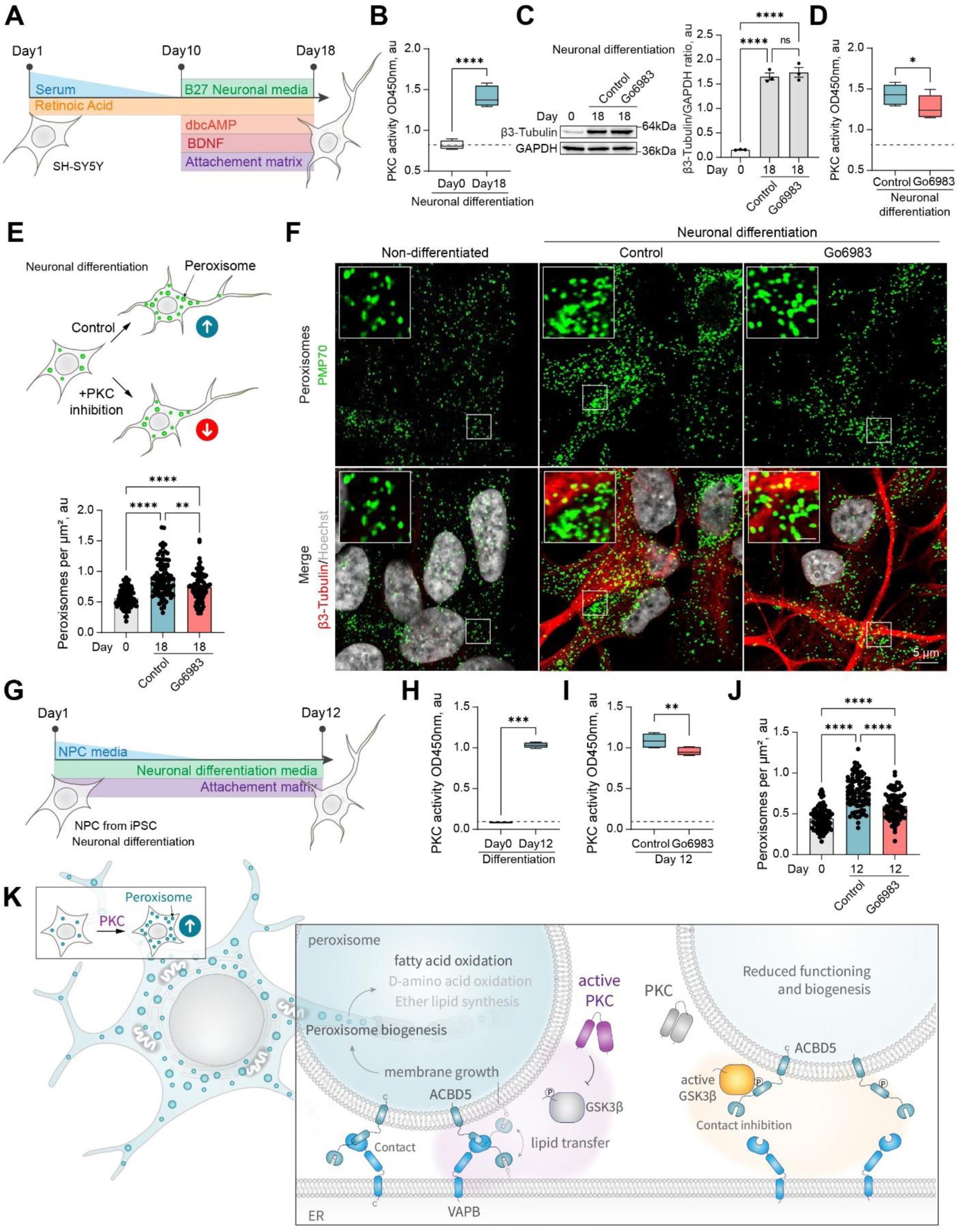
Protein kinase C regulates peroxisome abundance in neurons **(A)** Schematic of the SH-SY5Y neuronal differentiation. **(B)** PKC activity in the non-differentiated and 18-day differentiated SH-SY5Y cells. Quantification shows PKC activity, mean± SEM, N=10, **** - p<0.0001. **(C)** Western blot of β-3 tubulin in the non-differentiated and 18-day differentiated SH-SY5Y cells in control and Go6983 1μM conditions, Go6983 was added on days 10-18 of the differentiation protocol. Quantification shows the ratio of and β-3 tubulin to GAPDH, mean± SEM, N=3, **** - p<0.0001. **(D)** PKC activity in the 18-day differentiated SH-SY5Y cells in control and Go6983 1μM conditions, Go6983 was added on days 10-18 of the differentiation protocol. Quantification shows PKC activity, mean± SEM, N=10, * - p<0.05. **(E-F)** Confocal microscopy of peroxisomes in the non-differentiated and 18-day differentiated SH-SY5Y cells in control and Go6983 1μM conditions, Go6983 was added on days 10-18 of the differentiation protocol. Peroxisomes were visualized with the PMP70 antibody, nuclei were stained with Hoechst (10μg/ml), and neuronal differentiation was visualized with β-3 tubulin antibody. Representative images are shown, Scale bar - 5μm, inlet - 1μm. (F) Quantification shows the number of peroxisomes per square micron of the cytoplasm in the 2D confocal image, mean± SEM, N=102, **** - p<0.0001. Refer to Figure S12C. **(G)** Schematic of the neuronal progenitor cells (NPCs) neuronal differentiation. **(H)** PKC activity in the non-differentiated and 12-day differentiated NPCs. Quantification shows PKC activity, mean± SEM, N=8, **** - p<0.0001. **(I)** PKC activity in the 12-day differentiated NPCs in control and Go6983 1μM conditions, Go6983 was added on days 1-12 of the differentiation protocol. Quantification shows PKC activity, mean± SEM, N=8, ** - p<0.01. **(J)** Quantification of the number of peroxisomes per square micron of the cytoplasm in the 2D confocal image in the non-differentiated and 12-day differentiated NPCs in control and Go6983 1μM conditions, Go6983 was added on days 1-12 of the differentiation protocol, mean± SEM, N=90, **** - p<0.0001. Refer to Figure S12E. **(K)** Model of PKC regulation of peroxisome abundance

## Discussion

We established a mechanistic connection between PKC activation and peroxisome biogenesis through PEX11b peroxisome formation. We showed that PKC promotes VAPB-dependent peroxisome biogenesis, through its established role in GSK3β inhibition and, a recently discovered, GSK3β regulation of peroxisome-ER interaction^68^. This mechanism allows cells to rapidly induce peroxisomes through signaling regulation, providing both an on-demand pool of functional peroxisomes and fulfilling the need for peroxisomal function during neuronal differentiation (Figure 5K).

PKC is a Serine/Threonine kinase that controls cell-to-cell recognition, cytoskeletal dynamics, GSK3 activity, membrane growth, eisosome assembly, T-cell development, apoptosis, response to oxidative stress, and lysosome biogenesis, among many other cellular functions ^75, 76, 98, 103–114^. The PKC superfamily consists of 4 distinct sub-groups ^75, 76^ that share a kinase domain, and have a variety of regulatory domains that regulate spatiotemporal specificity ^75^ (Figure 2A). A screen of kinase inhibitor compounds revealed that inhibition of cPKC and nPKC results in a reduction of peroxisome numbers in human cell lines, including neuron-like cells. Peroxisome abundance is a balance between several pathways that produce and degrade peroxisomes, as well as PPARalpha-regulated transcriptional response that produces PEX peroxisome biogenesis factors ^38, 46, 82^. Classical isoforms of PKC regulate PPAR alpha receptor ^115^. One possibility is that PKC inhibition leads to reduced transcription of PEX genes, however we didn’t see an effect of PKC inhibition on the PPAR transcriptional response. Another possibility is that PKC inhibition increases pexophagy. However, blocking pexophagy in normal conditions in human cells did not lead to a significant change in peroxisome number. The regulation of peroxisome abundance by pexophagy may be more relevant for clearance of peroxisomes, when excess peroxisomes are degraded, usually after a bout of upregulation ^46, 81, 82, 89, 116^. The timing of PKC-induced peroxisome induction, as well as its dependence on PEX11b, suggested peroxisome division as the mechanism of PKC regulation. Our data showed that PKC positively regulates peroxisome-ER contact by modulating the ACBD5-VAPB tether by inhibiting GSK3β, which was recently shown to negatively regulate the contact^58, 68^. Although PKC can phosphorylate PEX11b on the N-terminus S53, the phosphorylation event is not sufficient to increase the number of peroxisomes. We also didn’t observe any effect of PKC on PEX11b interaction with other division factors. It remains to be determined how peroxisome division depends on peroxisome-ER contact sites ^117^.

The PKC-regulated pathway that we have identified has broad implications for peroxisome physiology. For example, PKC and peroxisomes are known regulators of redox signaling, and peroxisomes both produce and clear hydrogen peroxide ^118–120^. Both PKC and peroxisomes are known to control levels of D-serine in the brain ^121–125^. The model of PKC-induced peroxisome proliferation also explains the notable increase in peroxisome abundance during neuronal differentiation. PKC activity has been recorded to be the highest in the brain, followed by liver and kidney ^101^, the organs which incidentally are primarily affected in peroxisome biogenesis disorders (Peroxisome Biogenesis Disorder/Zellweger syndrome, also known as cerebro-hepato-renal syndrome) ^32, 126^. It would be of interest to explore the peroxisomal contribution to diseases associated with abnormal PKC signaling, such as cancer, cardiovascular diseases, diabetes, psoriasis, and neurodegenerative disorders ^127–133^.

## Acknowledgments

We thank Prof. Gisou van der Goot and Prof. Giovanni D’Angelo and members of their labs for their support, including access to essential equipment, reagents, and discussions. We thank Laurence Gouzi Abrami and Sylvia Ho for technical assistance. We thank the Gene Expression Core Facility (GECF, EPFL) for providing access to essential equipment and lenti-viral collection, Dr. Adrien Schmid and the Proteomics Core Facility (PCF) at EPFL for phospho-proteomic analysis, and the Bioimaging and Optics Core facility (PTBIOP, EPFL) for access to essential imaging equipment. We thank the Imaging and Microscopy centre (IMC) facility, University of Southampton. We thank Dr. Julien Schmidt and the Peptide and Tetramer Core Facility, University of Lausanne for peptide synthesis. We thank Prof. Ronald J. A. Wanders for the advice on radioactive measurements of peroxisomal fatty acid oxidation. We thank Prof. Doug Kellogg for a generous gift of the Pkc1 antibody. T.A. was funded by the HFSP Long-term Fellowship (LT000559/2021-L), EPFL Faculty, the University of Southampton, and the Wessex Medical Research Innovation Grant. A.B. was funded by the HFSPO Scientists for Scientists Initiative to help scientists affected by the war in Ukraine.

## Author contributions

A. B., C. H., S. P. and T. A. performed the experiments. T. A. and D. K. designed the experiments, performed the data analysis, wrote the manuscript, and supervised the study.

## Methods

### Materials Availability

Reagents generated in this study are available upon request.

### Data and Code Availability

Additional data that support the conclusions of this study are available upon request. This study didn’t generate any code.

## EXPERIMENTAL MODEL AND SUBJECT DETAILS

Human fibroblasts (AFF11), U2OS, HeLa, CHO, and HEK293T cells were maintained in high glucose DMEM supplemented with 10% fetal bovine serum (FBS), 1% penicillin/streptomycin, at 37°C/5% CO2, SH-SY5Y cells were maintained in high glucose DMEM/F12 1:1 media supplemented with 10% FBS, 1% penicillin/streptomycin at 37°C/5% CO2. Cells modified via CRISPR/Cas9 were maintained as above with addition of puromycin (1μg/ml, Sigma) or Blasticidin (5μg/ml, Sigma) during selection of the clonal populations. Neural progenitor cells were maintained in STEMdiff™ neural progenitor medium (STEM CELL Technologies). Neuronal differentiation of SH-SY5Y cells was done according to ^102^ with modifications, SH-SY5Y cells (initial seeding density 350,000 cells on 60mm dish) were gradually FBS starved for 10 days, before plating them on a Matrigel (Corning) extra-cellular matrix (seeding density (100,000 per a well of a 6-well plate), neuronal media was replaced every third day, cells were collected on day 18. Neuronal differentiation of induced pluripotent stem cells (iPSC) derived neuronal progenitor cells (NPC) ^134^ was done using STEMdiff™ neural differentiation kit (STEM CELL Technologies). The concentration of cells for plating was determined using cell counter (Countess II FL, Life Technologies) with the cell counting chambers (Invitrogen).

## METHOD DETAILS

### Antibodies

We used the following reagents to detect proteins: anti-GAPDH (sc-47724, Santa Cruz Biotechnology), anti-mCherry (34974, Invitrogen), anti-PMP70 (SAB4200181, Sigma), anti-PEX19 (14713-1-AP, Proteintech), anti-GFP (SAB4301138, Sigma), anti-NBR1 (16004-1-AP, Proteintech), anti-PEX11b (PA5-37011, Thermo Fisher Scientific), anti-PKC (P5704, Sigma), anti-Flag (F1804, Sigma), anti-myc (9E10, Sigma), anti-MFF (17090-1-aP, Proteintech), β-3 tubulin (D71G9, Cell Signaling Technology), anti-VAPB (14477-1-AP, ProteinTech).

Secondary antibodies for immunofluorescence: anti-Rabbit IgG Cy3-conjugated (Sigma-Aldrich C2306), anti-Mouse IgG Cy3-conjugated (Sigma-Aldrich C2181), anti-rabbit IgG Cy5 conjugated (Invitrogen A10523), Anti-Mouse IgG H&L (Alexa Fluor® 488) (Abcam).

### Chemicals

Hoechst (Sigma), fatty acid free BSA (PAN), Phenylmethylsulfonyl fluoride (PMSF, Sigma), Go6983 (Cayman Chemicals), Behenic acid (VLCFA, Sigma), kinase screening library (Cayman Chemicals), D-Serine (Sigma), palmitic acid (LCFA, Sigma), Phorbol 12-myristate 13-acetate (Cayman Chemicals), siRNA NBR1 and control (Qiagen), oleic acid (Sigma), brain derived neurotrophic factor (BDNF, Stem Cell Technologies), Dibutyryl-cAMP (Santa Cruz Biotechnology), Neuropan 27 Supplement 50x (Pan Biotech).

### CRISPR/Cas9

Knockout and endogenously tagged cell lines were constructed using CRISPR/Cas9 protocol and plasmids described in Ran et. al (Ran et al., 2013). Knockout cell lines were verified by western blotting, immunofluorescence. Genomic DNA was sequenced to verify disrupted region in knockout or the fidelity of endogenous tagging. Functional assay for knockout verification was performed where applicable (Reduced viability of PEX19 on VLCFA). Endogenous tagging was performed by fusing tagging construct (linker-GFP-polyA-Blasticidinn for C-terminal tagging of PMP70, and flag- GFP- for the N-terminal tagging of PEX11b) to the region upstream of the stop codon (∼500bp) and downstream of stop codon (∼500bp) for the C-terminal tagging, and to the region upstream of the start codon, and downstream of the start codon for the N-terminal tagging ^135^. PCR product containing homologous regions flanking the tagging construct was co-transfected with px330-gRNA corresponding construct for C-terminal tagging and px459-gRNA construct for N-terminal tagging. Endogenous tagging was verified by western blotting, immunofluorescence staining, and genomic DNA sequencing. CRISPR specificity was profiled using Digenome-Seq web tool (http://www.rgenome.net/cas-offinder/) (Bae et al., 2014). Off targets were not found. The following target sequences are used to modify genomic DNA: endogenous tagging of PMP70 on the C-terminus – 5’-GTTGAGTTTGGCTCTTAGAGAAATC-3’, endogenous tagging of PEX11b on the N-terminus – 5’-CGCGGAGCCTGGGCTGCGGCTGTCA-3’, knockout of PEX19 – 5’-GGAGGTAGCAAGATGGCCGCCGCTG-3’, knockout of PEX11b - 5’-CGCGGAGCCTGGGCTGCGGCTGTCA-3’.

### Plasmid Construction

All plasmids were constructed using *Escherichia coli* strain DH5α. Plasmids used in this study are summarized in the Table 1. We used px459 and px330 plasmids to clone CRISPR/Cas9 constructs for gene knockout and endogenous tagging. pSpCas9(BB)-2A-Puro (PX459) V2.0 was a gift from Feng Zhang (Addgene plasmid # 62988 ; http://n2t.net/addgene:62988; RRID:Addgene_62988) (Ran et al., 2013). pX330-U6-Chimeric_BB-CBh-hSpCas9 was a gift from Feng Zhang (Addgene plasmid # 42230 ; http://n2t.net/addgene:42230; RRID:Addgene_42230) (Cong et al., 2013). pcDNA4-PKCZeta WT His tagged was a gift from Jeff Wrana (Addgene plasmid # 24609 ; http://n2t.net/addgene:24609; RRID:Addgene_24609) ^136^. PKC alpha WT was a gift from Bernard Weinstein (Addgene plasmid # 21232 ; http://n2t.net/addgene:21232; RRID:Addgene_21232) ^137^. pWZL Neo Myr Flag PRKCD was a gift from William Hahn & Jean Zhao (Addgene plasmid # 20603 ; http://n2t.net/addgene:20603; RRID:Addgene_20603) ^138^. HA GSK3 beta wt pcDNA3 was a gift from Jim Woodgett (Addgene plasmid # 14753 ; http://n2t.net/addgene:14753; RRID:Addgene_14753)^139^. pCDNA3.1-PPARA was a gift from Claes Wadelius (Addgene plasmid # 169019 ; http://n2t.net/addgene:169019; RRID:Addgene_169019)^140^. Human PEX3, PEX19, DAO, and ACBD5 were amplified from Lentiviral plasmid collection. Side Directed Mutagenesis was verified by sequencing. Site directed mutagenesis was performed to obtain PEX11b mutants, and GSK3bS9A.

**Table 1.** Plasmids used in this study.

| Plasmid name | Source |
| --- | --- |
| pcDNA3.1 mCH*4skl | <sup>134</sup> |
| pcDNA3.1 GSK3beta | Addgene 14753 |
| pcDNA3.1 PPARA | Addgene 169019 |
| pcDNA3.1 PEX3hismyc | This study |
| pcDNA3.1 flagPEX19 | This study |
| Px459-PEX19 KO | <sup>66</sup> |
| Px330-PMP70-C | This study |
| Px459-PEX11b-N | This study |
| pCtag-PMP70-GFP-Blasti | This study |
| pNtag-PEX11b-flag-GFP | This study |
| pcDNA3.1- PKCα-mCherry | This study, PKCα is from Addgene 21232 |
| pcDNA3.1 PKCδ -mCherry | This study, PKCδ is from Addgene 20603 |
| pcDNA3.1- PKCζ -mCherry | This study, PKCζ is from Addgene 24609 |
| pcDNA3.1 hismyc-PEX11b | This study |
| pcDNA3.1 hismyc-PEX11b S53D | This study |
| pcDNA3.1 hismyc-PEX11b S53A | This study |
| pEGFP-VAPB | <sup>141</sup> |
| pcDNA3.1 hismyc-ACBD5 | This study |
| pcDNA3.1 GFP-SKL | This study |
| pcDNA3.1 ER-mCherry | <sup>134</sup> |

### PKC activity

Protein kinase activity was determined by PKC Kinase Activity Assay Kit (Abcam) according to the manufacturer instructions. Protein concentration in lysates was quantified using bicinchoninic acid (BCA) Assay kit (Interchim).

### Peroxisome abundance measurement

We used peroxisome density as a read-out for peroxisome abundance in this study. To calculate the number of peroxisomes cells were visualized by confocal microscopy. Images were captured using the same parameters between conditions within the experiment. Images were analyzed by Fiji (ImageJ) software, peroxisome numbers were quantified as maxima of intensity in the cytoplasmic space and the number was divided by the size of the areas (square micron) which results in a peroxisome per square micron cytoplasmic ratio. The cytoplasmic space was defined using threshold mask that excludes the nuclei (stained with Hoechst) and background using ImageJ Threshold function. The resultant area was divided into N parts equal to the number of cells (nuclei number). At least 100 areas were quantified per condition per experiment. 2D areas with a nucleus in focus were chosen for all of the experiments, because 3D projections often create overlaps between peroxisomal compartments.

### Preparing samples for the Mass spectroscopy

Cells expressing CRISPR/Cas9 flag-GFP-PEX11b were grown to 80% confluency in the presence of VLCFA for 2 days, washed with PBS, and lysed in 1%TritonX-100 50mM Tris HCl pH7.4, 150mM NaCl buffer at 4°C for 30min. PEX11b was purified using Anti-FLAG M2 affinity gel (Sigma). 10% of the sample was collected for the Western Blot analysis. Beads were collected and washed free of detergents in 50mM Tris HCl, pH7.4 150mM NaCl. Samples were reduced in 10 mM dithioerythritol (DTE) alkylated in 55 mM iodoacetamide (IAA)). Digestion was performed overnight at 37 °C using mass spectrometry grade trypsin and LysC protease (Thermo Fisher Scientific). The following day, protein digests were then subjected to C18 stage tip cleaning, dried in a speed-vacuum and stored at - 20°C until LC-MS analysis.

### LC/MS/MS analysis (shotgun approach)

Mass spectrometry-based proteomics-related experiments were performed by the Proteomics Core Facility at EPFL. Mass spectrometry (MS) analysis was performed on an Orbitrap Exploris 480 mass spectrometer (Thermo Fisher Scientific) coupled to a nano-UPLC Dionex pump. For liquid chromatography (LC) MS/MS analysis, trypsin / LysC digested samples were resuspended in 30-60ul of a mobile phase A solution (2% ACN / water, 0.1% formic acic) and then separated by reversed-phase chromatography using a Dionex Ultimate 3000 RSLC nanoUPLC system on a home-made 75 µm ID × 50 cm C18 capillary column (Reprosil-Pur AQ 120 Å, 1.9 µm) in-line connected with the MS instrument. Peptides were separated by applying a non-linear 90min gradient ranging from 99% solvent A (2% ACN and 0.1% FA) to 90% solvent B (90% ACN and 0.1%FA) at a flow rate of 250nl/min. For spectral library and charge state determination of Pex11b peptides, the MS instrument was operated in data-dependent mode (DDA). Full-scan MS spectra (300−1500 m/z) were acquired at a resolution of 120’000 at 200 m/z. Data-dependent MS/MS spectra were recorded followed by HCD (higher-energy collision dissociation) fragmentation on the ten most intense signals per cycle (2 s), using an isolation window of 1.4 m/z. HCD spectra were acquired at a resolution of 60’000 using a normalized collision energy of 32 and a maximum injection time of 100 ms. The automatic gain control (AGC) was set to 100’000 ions. Charge state screening was enabled such that unassigned and charge states higher than six were rejected. Precurors intensity threshold was set at 5’000. Precursor masses previously selected for MS/MS acquisition were excluded from further selection for a duration of 20 s, and the mass exclusion window was set at 10 ppm.

### MS data processing and database searches

PEAKS Studio X+ Pro (Bioinformatics Solutions Inc.) software was used for data processing. The raw MS data files were imported into PEAKS Studio software using the following parameters for the database search: For protein identification, the UniProt/Swiss-Prot human proteome data base (UP000005640) or searched using the canonical protein sequence of human Pex11b (accession#: O96011) combined with a decoy and contaminant database was used. For peptide identification the following settings were used: Enzyme: Trypsin, missed cleavages: 2 precursor mass tolerance: 10 ppm, fragment mass tolerance: 0.1 Da, minimum charge: 2, maximum charge: 5, fixed modifications: Carbamidomethyl (C), variable modifications: Oxidation (M), phosphorylation (STY). False discovery rate (FDR) was calculated based on the target/decoy database and peptides as well as proteins with FDR threshold of ≤ 1% (or -log10P ≥15 for peptides) were chosen as true positive hits. The PTM identification threshold (PEAKS) was set at the recommended AScore of -log10P of 20 (p=0.01).

### RNA preparation and real time PCR (rtPCR)

Total mRNA was extracted from cells using RNeasy Mini Kit (Qiagen). cDNA synthesis was performed using iScript cDNA synthesis kit (Biorad). Real time PCR was performed using QuantStudio6 (Thermo Fischer Scientific). mRNA levels were quantified using QuantStudio6 software. Experiments were repeated three times with 2 technical repeats and fold difference in expression was calculated by ΔΔCt method using GAPDH as a housekeeping gene (Livak and Schmittgen, 2001).

**Table 2:**
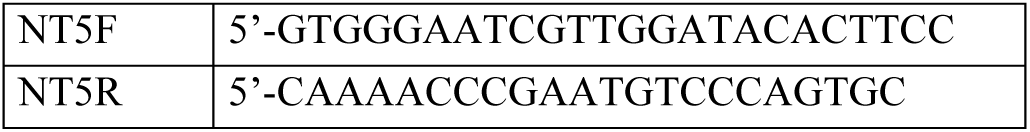
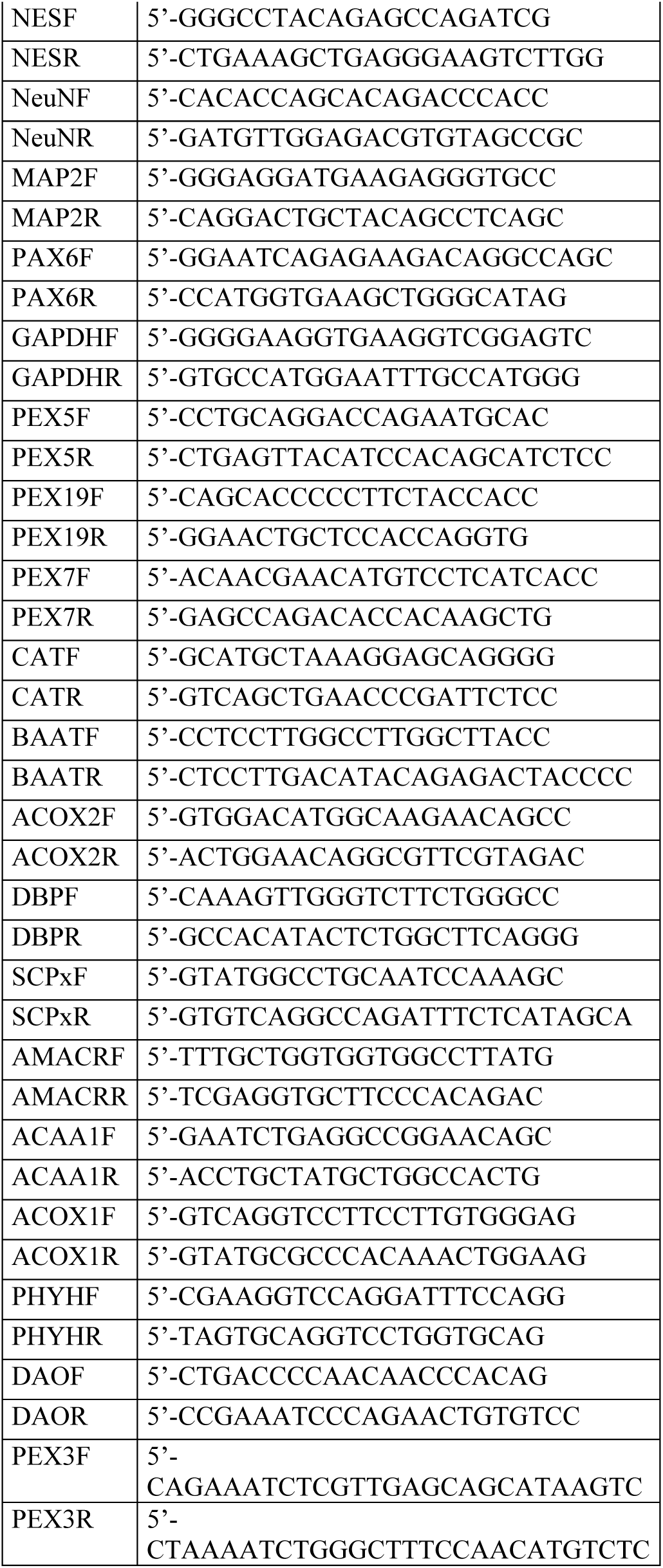
Primers for rtPCR.

### Co-Immunoprecipitation

Harvested cells were lysed with CHAPS (1%) in 50mM Tris-HCl pH7.4, 150mM NaCl with a complete protease inhibitor cocktail (Pierce) for 20 min at 4°C with rotation (1%CHAPS in PBS for ACBD5 immunoprecipitation). After 1 min 10,000g centrifugation the supernatant was incubated with protein G Sepharose (GE Healthcare) for 30min at 4°C with rotation. After 1min 10,000g centrifugation 10% of the supernatant was collected as an Input, and the resulting supernatant was incubated with 25μl of Myc-Trap Agarose beads (Chromotek) overnight at 4°C with rotation. After the incubation the beads were washed 4 times for 5 min with the lysis buffer at 4°C with rotation. Before loading on the gel, the beads and input were boiled with a sample buffer. Equal amounts of Input and Eluted bead sample were loaded on the SDS-PAGE.

### Immunofluorescence

Cells were grown on glass bottom plates or glass slides, fixed using 4% paraformaldehyde in phosphate buffered saline (PBS) for 10 minutes, washed with PBS, permeabilized with 0.5% Triton X100, then blocked overnight in 5% BSA in PBS prior to antibody staining.

### Radioactive measurement of beta fatty acid oxidation using 13,14-3H docosanoic acid (C22:0)

Cells were grown in 12-well plates with initial plating density of 100 000 cells per well for two days. Cold C22:0 (4 µM) and 13,14-3H C22:0 (1 µCi, Anawa) were mixed with the media and added to the cells for 20 hours. Media was collected and processed according to ^142, 143^ with minor modifications. Radioactivity was quantified in the media and water fractions transferred to the Scintillation vials after 3 days using scintillation counter. Cold samples, and cell-free samples were used as controls and background subtraction. Peroxisome deficient cells were used as a control to distinguish between peroxisomal and mitochondrial fatty acid oxidation.

### Microscopy

For live cell imaging 4-well microscope glass bottom plates (IBIDI) or Cellview cell culture dish (Greiner Bio One) were used. Alternatively, cells were grown on glass slides (Marienfeld). Confocal images and movies were acquired using SP8 (Leica) confocal microscope equipped with a temperature and CO2 incubator, using a 60x PlanApo VC oil objective NA 1.40. We used 406nm, 488nm, 561nm, and 640nm lasers. Peroxisome tracking in the movies was done using TrackMate Fiji plugin^144^. Super resolution microscopy was done using STED microscope (Leica TCS SP8 STED 3X) with a 100 × 1.4 NA oil-immersion objective. Image processing was performed using Fiji (ImageJ) software.

### Peptide phosphorylation competition assay

Peptides were synthesized by the Peptide and Tetramer core Facility, University of Lausanne. The following peptides were used for the assay: PEX11b-1 GCGGLESHLSLGRKL, PEX11b-2 GCGGADALESAKRAV, PEX11b-3 GCGGKWAQRSFRYYL, PEX11b-4 GCGGRYYLFSLIMNL, PEX11b-5 GCGGESSACSRRLKG;PEX11b-6 GCGGRRLKGSGGGVP, PKCa-Control GCGGFKKQGSFAKKK ^95^. 100ng of pure peptides were mixed with or without 100ng of pure PKC, and incubated according to the PKC activity assay kit (Abcam). Peptides were then subjected to mass spectrometry.

## QUANTIFICATION AND STATISTICAL ANALYSIS

Three or more independent experiments were performed to obtain the data. P values were calculated by two-tailed Student t-test, or one-way ANOVA for samples following normal distribution determined by the Shapiro-Wilks test. The equality of variances was verified by Brown-Forsythe or F test. Mann-Whitney (2 groups), or Kruskal-Wallis (multiple groups) tests were used for samples that didn’t follow a normal distribution. The sample sizes were not predetermined. Refer to the Supplementary Table 3 for the details on statistical analysis.

### Supplemental Items

Table S1. Kinase Inhibitor Screen

Table S2. Phosphorylation of PEX11b in VLCFA conditions

Table S3. Statistical Analysis

Supplementary Movies 1-6

Supplementary Figures S1-S10

---

**Figure S1.**
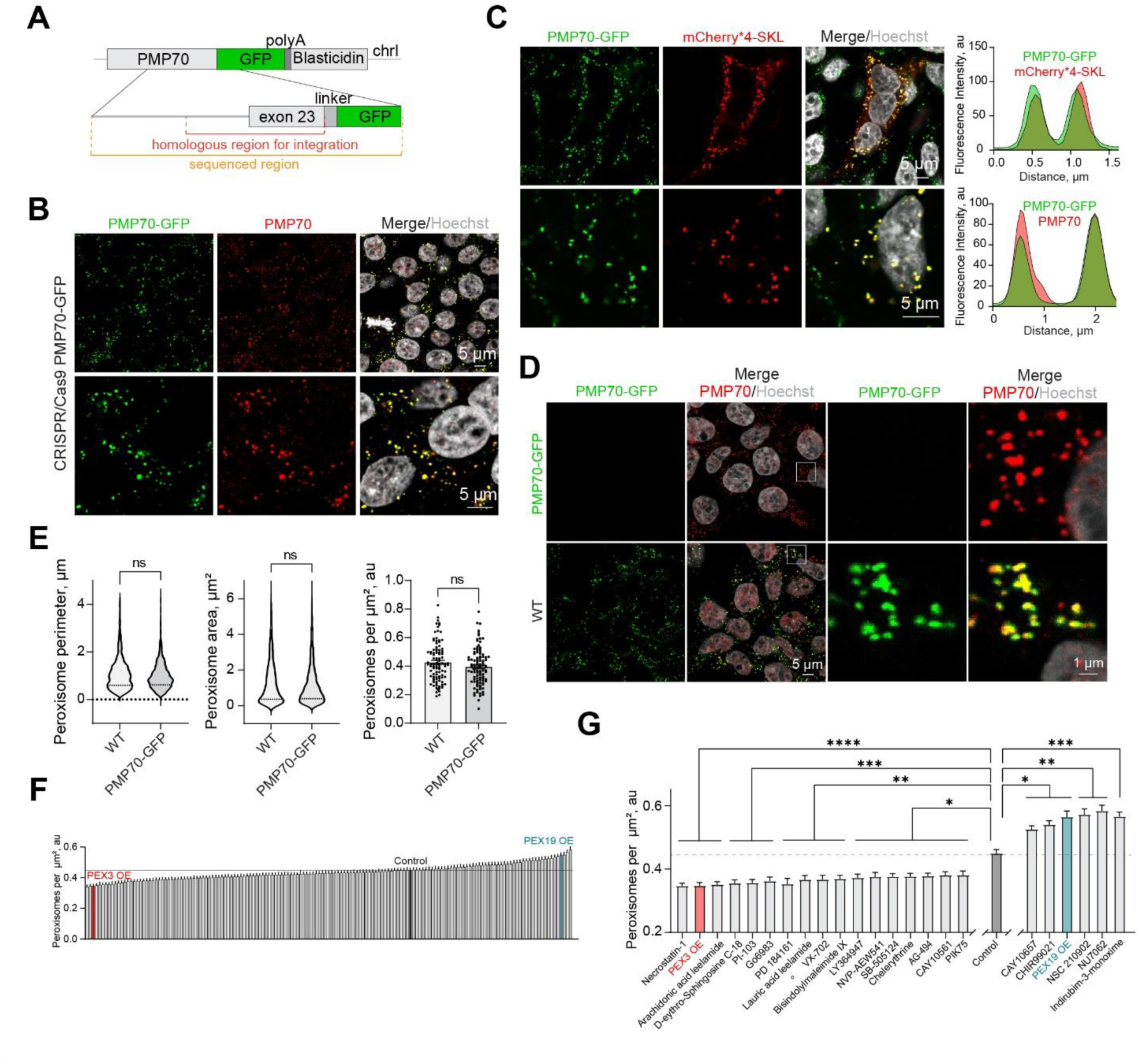
**(A)** Schematic of the genomic DNA region corresponding to the end of the human PMP70 open reading frame endogenously tagged with -GFP-polyA-Blasticidin. **(B-C)** Confocal microscopy of the peroxisomal import marker mCherry*4-SKL expressed in HEK293T PMP70-GFP, and (**B**) a confirmation of the PMP70-GFP peroxisomal localization. The fluorescence intensity profiles through single peroxisomes are shown. Scale bar -5μm. **(D-E)** Comparison of WT and PMP70-GFP tagged HEK293T. Confocal images and quantification of perimeter (N=1000), area (N=1000), and density of peroxisomes (N=100) are shown, mean± SEM. **(F-G)** Quantification of the number of peroxisomes per square micron of the cytoplasm in the HEK293T cells treated with 1μM of the indicated small molecule, mean± SEM, * - p<0.05, ** - p<0.01, *** - p<0.001, **** - p<0.0001. **(D)** – significant screen hits are plotted, **(E)** complete screen. Refer to Figure S2 and S3 for the confocal images, N=100 for each molecule.

**Figure S2.**
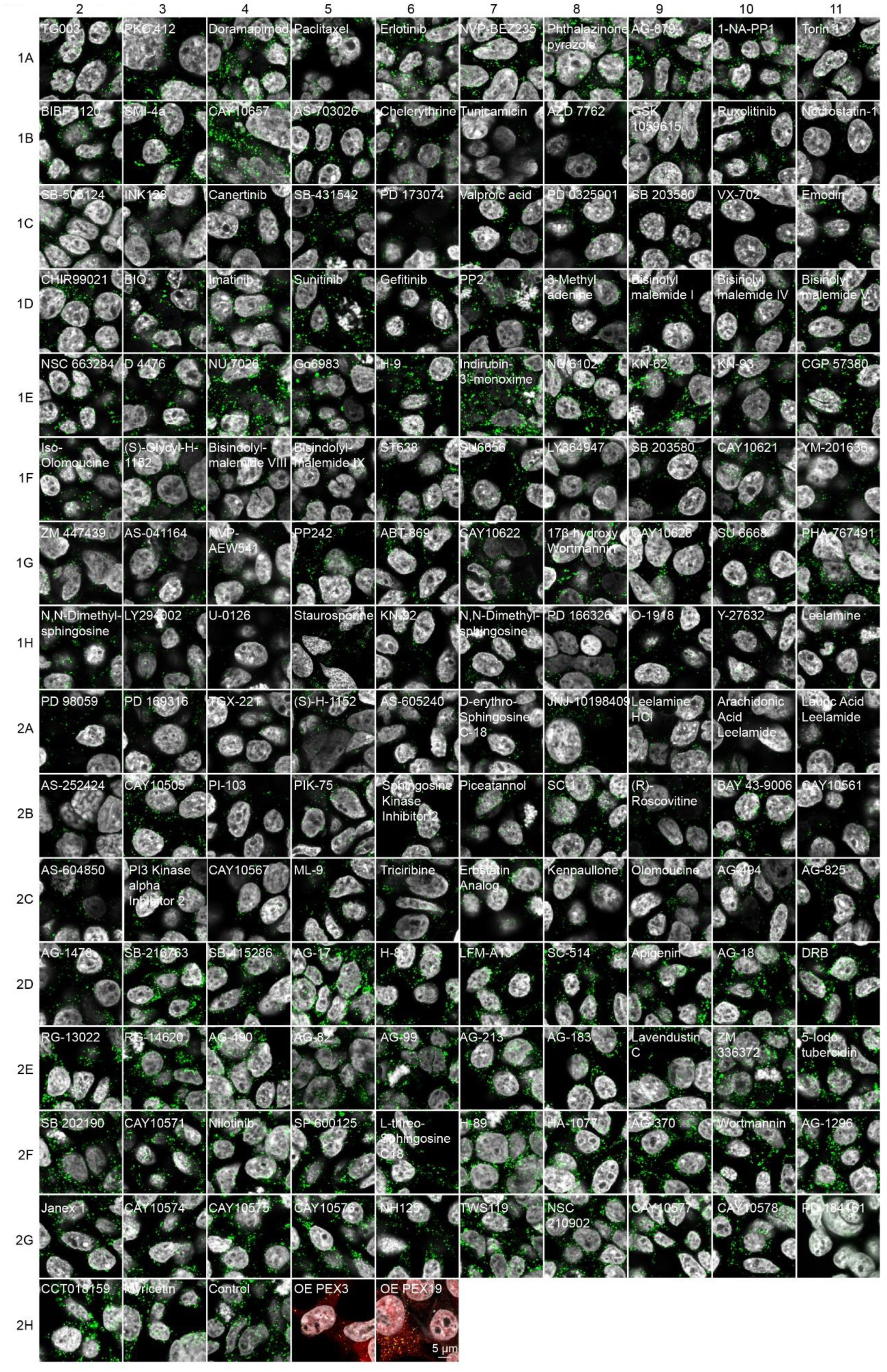
Kinase Inhibitor Screen. Confocal microscopy of the HEK293T CRISPR/Cas9 PMP70-GFP cells incubated for 2 days with 1μM of the indicated small molecule. Note, that endogenous levels of PMP70 vary between treatments as do the numbers of peroxisomes. Scale bar - 5μm.

**Figure S3.**
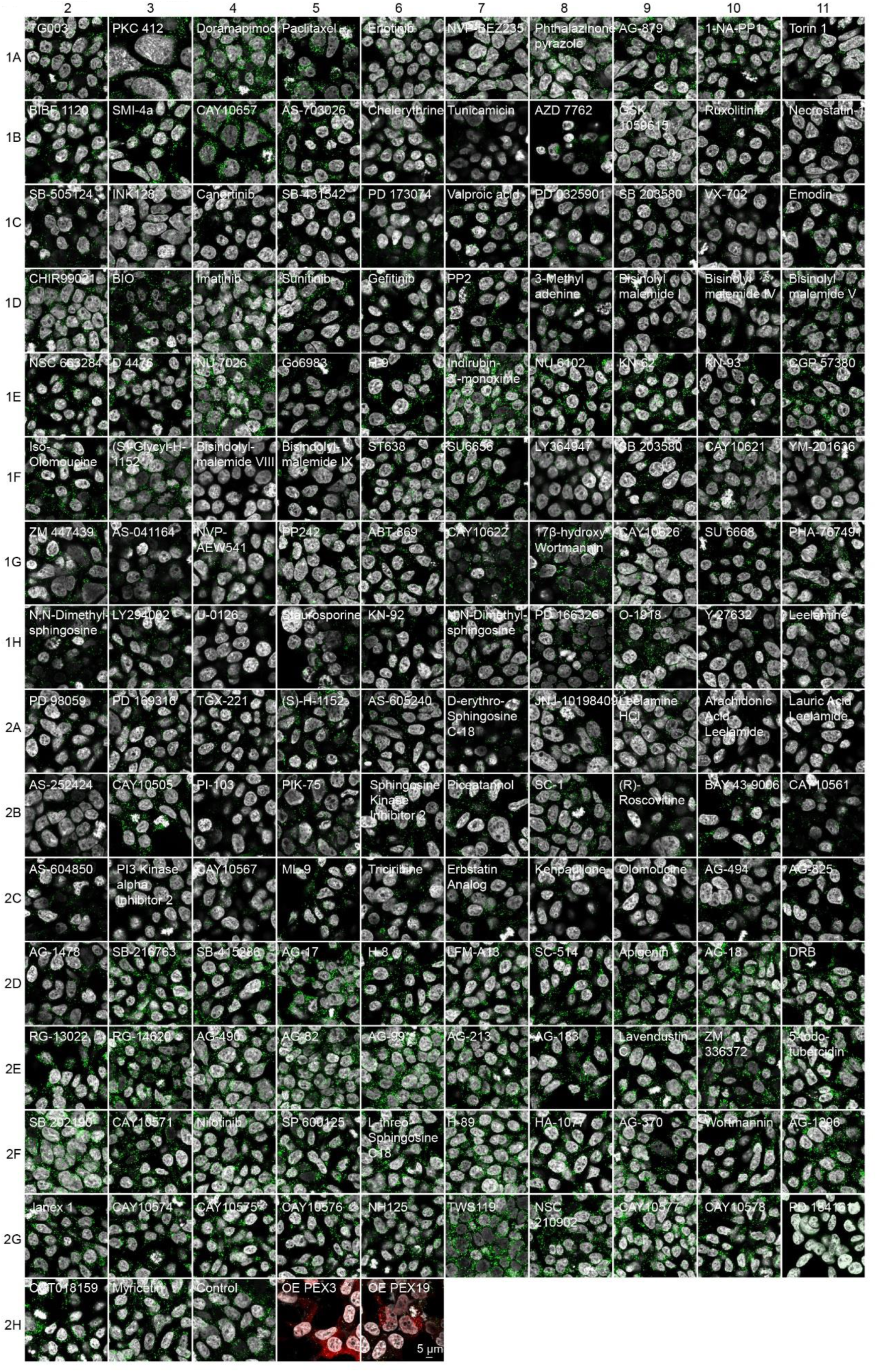
Kinase Inhibitor Screen. Confocal microscopy of the HEK293T CRISPR/Cas9 PMP70-GFP cells incubated for 2 days with 1μM of the indicated small molecule. Note, that endogenous levels of PMP70 vary between treatments as do the numbers of peroxisomes. Scale bar - 5μm.

**Figure S4.**
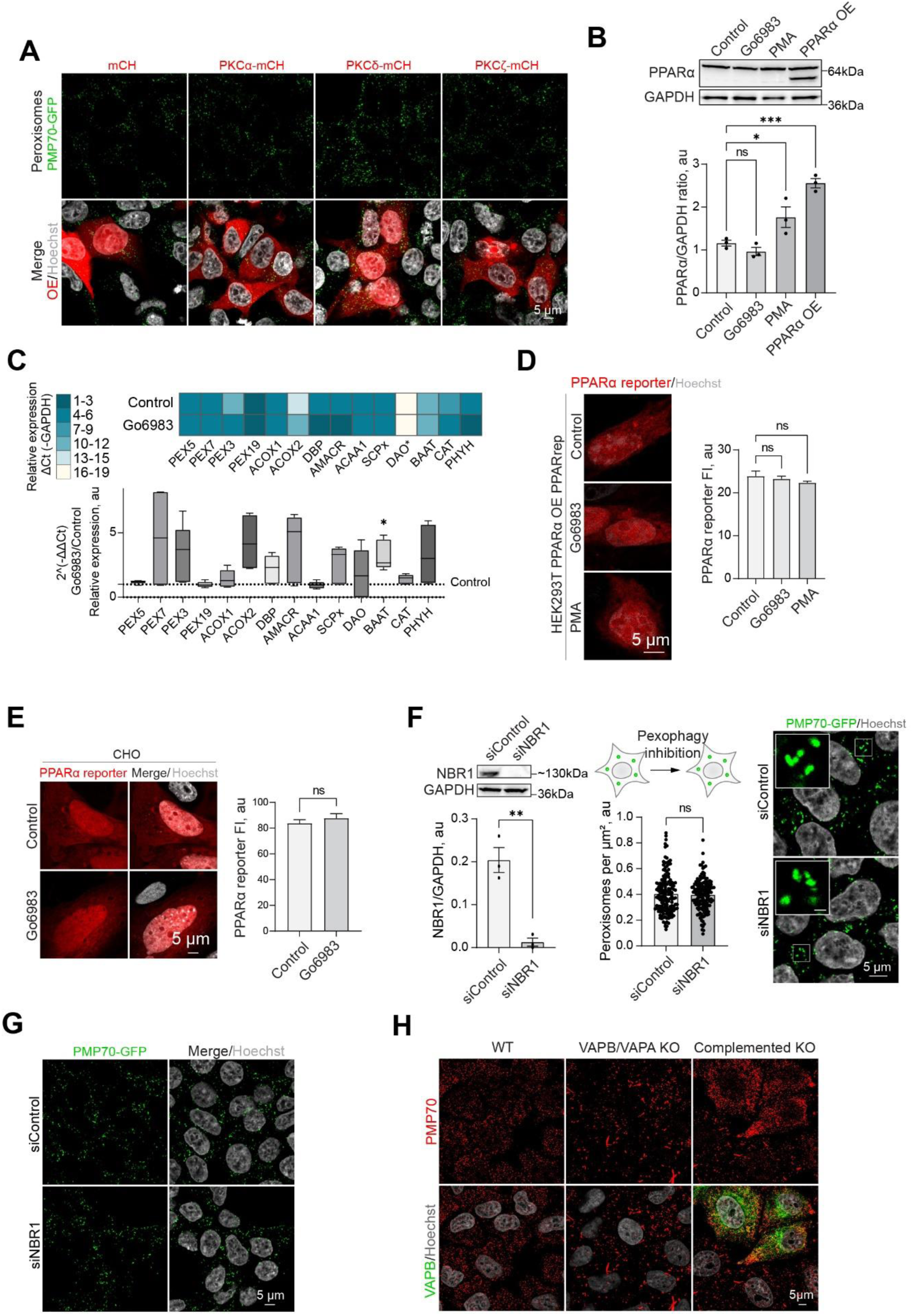
**(A)** Confocal microscopy of HEK293T CRISPR/Cas9 PMP70-GFP cells overexpressing (OE) PKCα-mCherry, PKCδ-mCherry, or PKCζ-mCherry. Scale bar - 5μm. **(B)** Western blot of PPARα in HEK293T cells in control PMA (0.5μM for 1day) and Go6983 (5μM for 2days) conditions. Quantification shows the ratio of PPARα to loading control, mean± SEM, N=3, *** - p<0.001, * - p<0.05 **(C)** Quantitative PCR of the peroxisomal genes in HEK293T cells in control and Go6983 5μM conditions (2 day treatment). Quantification shows color coded relative expression levels calculated as a ΔCt (Peroxisomal gene - GAPDH expression reference), and a ratio of Go6983/Contol expression, expressed as 2^-ΔΔCt^, mean± SEM, N=4, * - p<0.05. **(D-E)** Confocal microscopy of PPAR mCherry reporter in HEK293T cells expressing PPARα (D) and CHO (E) cells in control, Go6983 (5μM for 2days), or PMA conditions (0.5μM for 1day). Nuclei were stained with Hoechst (10μg/ml). Representative images are shown, Scale bar - 5μm. Quantification shows average fluorescence intensity per cell, N=50, **** - p<0.0001. **(F-G)** Western blot NBR1 and control silencing in HEK293T cells. Quantification shows the NBR1/GAPDH ratio quantified by the lane intensity on the western blot, mean± SEM, N=3, **** - p<0.0001. (G) Confocal microscopy of peroxisomes in HEK293T CRISPR/Cas9 PMP70-GFP cells in control silencing or NBR1 silencing conditions. Nuclei were stained with Hoechst (10μg/ml). Representative images are shown, Scale bar - 5μm, inlet - 1μm. Quantification shows the number of peroxisomes per square micron of the cytoplasm in the 2D confocal image, mean± SEM, N=150. **(H)** Confocal microscopy of peroxisomes stained with antiPMP70 antibody in WT and VAPB/VAPA KO HeLa cells in control and GFP-VAPB overexpression conditions.

**Figure S5.**
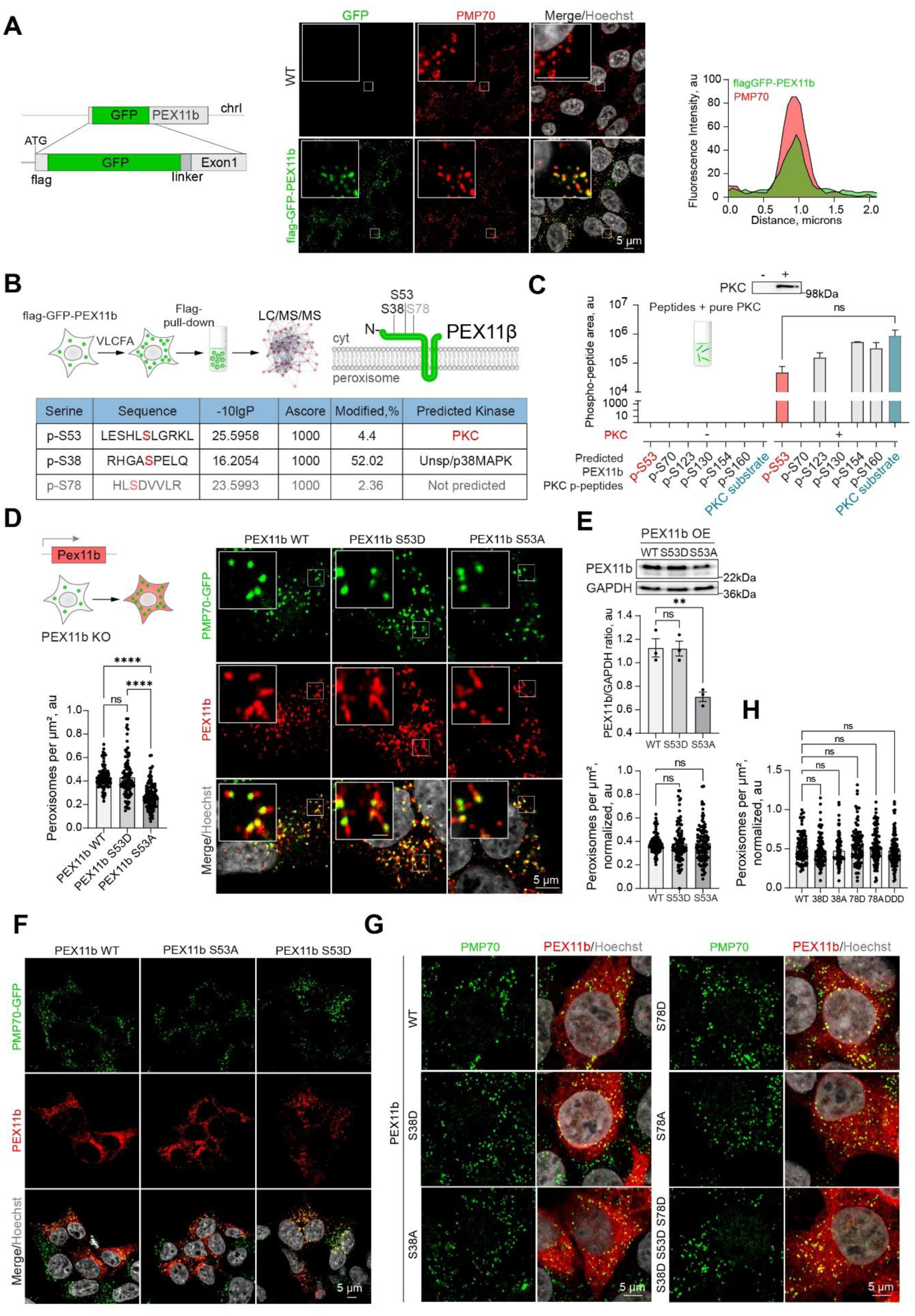
PKC phosphorylation of PEX11b on S53 is not sufficient to induce peroxisome proliferation **(A)** Confocal microscopy of flag-GFP-PEX11b CRISPR/Cas9 tagged HEK293T cells. Peroxisomes were visualized with PMP70 antibody, nuclei were stained with Hoechst (10μg/ml). Scale bar - 5μm, inlet - 1μm. Intensity profile through a single peroxisome is shown. Schematic of CRISPR/Cas9 flag-GFP-PEX11b construction, showing the N-terminal region of genomic DNA corresponding to PEX11b ORF. **(B)** Identification of PEX11b phosphorylation by targeted LC-MS/MS analysis, showing the identification of site-specific phosphorylation at residues S38, S53 and S78. **(C)** Phosphorylation of pure peptides (a mix of PEX11b peptides predicted to be phosphorylated by PKC, and a PKC-specific substrate peptide) incubated with or without active PKC. Western blot shows PKC in the treated and non-treated samples. Quantification shows the size of the peptide area identified by MS, mean± SEM, N=3, comparison between PKC-specific substrate and the PEX11b peptides. **(D)** Confocal microscopy of peroxisomes and PEX11b in HEK293T CRISPR/Cas9 PMP70-GFP PEX11b KO cells overexpressing myc-PEX11b, myc-PEX11b S53A, or myc-PEX11b S53D. Nuclei were stained with Hoechst (10μg/ml). Representative images are shown, Scale bar - 5μm, inlet - 1 μm. Quantification shows the number of peroxisomes per square micron of the cytoplasm in the 2D confocal image, N=100, mean± SEM. ****- p<0.0001. **(E)** Western blot of myc-PEX11b, myc-PEX11b S53A, or myc-PEX11b S53D, N=3, and corrected quantification showing the number of peroxisomes per square micron, mean± SEM, N=100. **(F)** Confocal microscopy of peroxisomes and PEX11b in HEK293T CRISPR/Cas9 PMP70-GFP PEX11b KO cells overexpressing myc-PEX11b or myc-PEX11b S53D. Nuclei were stained with Hoechst (10μg/ml). Representative images are shown, Scale bar - 5μm. **(G-H)** Confocal microscopy of peroxisomes and PEX11b in HEK293T overexpressing myc-PEX11b or myc-PEX11b indicated mutants. Nuclei were stained with Hoechst (10μg/ml). Representative images are shown, Scale bar - 5μm. Quantification shows the density of peroxisomes normalized by PEX11b expression level quantified by fluorescence intensity, mean± SEM, N=100.

**Figure S6.**
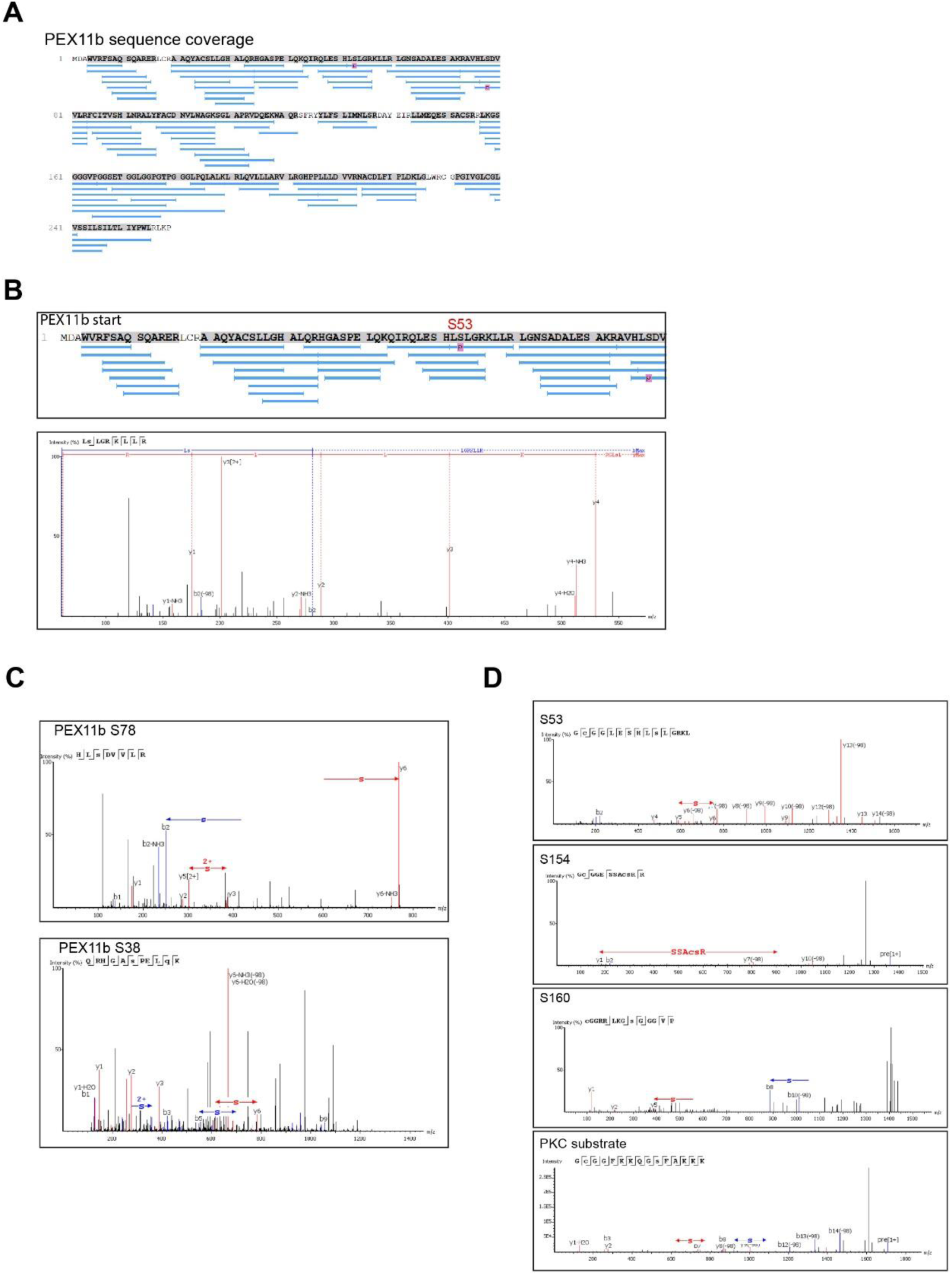
**(A)** Peptide coverage of targeted LC-MS/MS analysis of PEX11b **(B)** MS/MS spectrum from a targeted, LC-MS/MS analysis of Pex11b peptides, showing the identification of site-specific phosphorylation at residue S53. (Top) PEX11b sequence with identified peptide sequences (blue lines) and a site-specific phosphorylation. (Bottom) MS/MS spectrum of peptide the sequencing and identification of different -b & -y ions. **(C)** MS/MS spectrum from a targeted, LC-MS/MS analysis of Pex11b peptides, showing the identification of site-specific phosphorylation at residue S38, and S78. **(D)** Examples of MS/MS spectrums from a peptide competition assay, showing PEX11b phosphor-peptides as well as the PKC-specific phospho-substrate.

**Figure S7.**
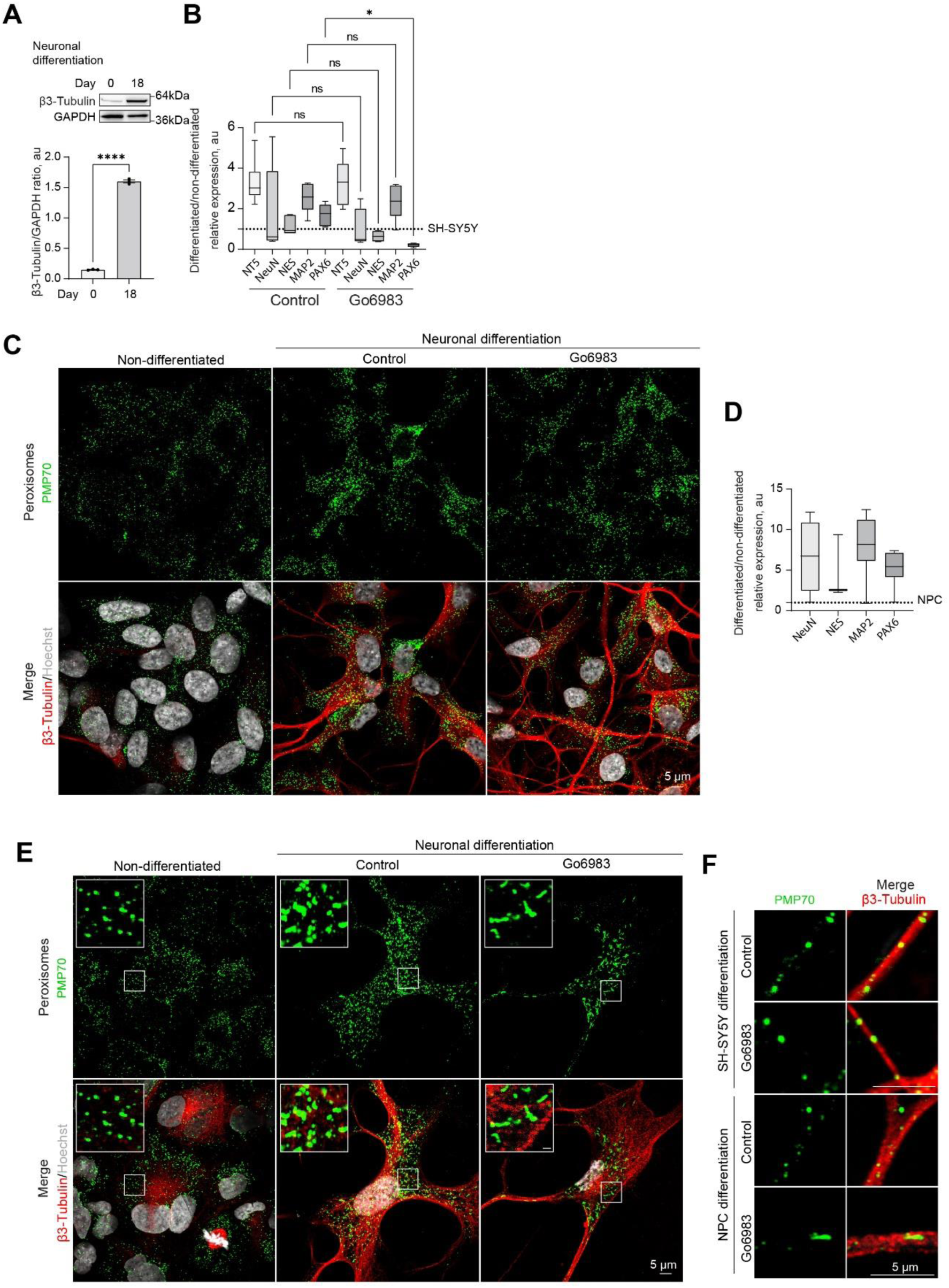
**(A)** Western blot of β-3 tubulin in the non-differentiated and 18-day differentiated SH-SY5Y. Quantification shows the ratio of and β-3 tubulin to GAPDH, mean± SEM, N=3, **** - p<0.0001. **(B)** Quantitative PCR of the neuronal markers in the 18-day differentiated SH-SY5Y cells in control and Go6983 1μM conditions, Go6983 was added on days 10-18 of the differentiation protocol. Quantification shows the relative expression of differentiated/non-differentiated markers, expressed as 2^-ΔΔCt^, mean± SEM, N=4, * - p<0.05. **(C)** Confocal microscopy of peroxisomes in the non-differentiated and 18-day differentiated SH-SY5Y cells in control and Go6983 1μM conditions, Go6983 was added on days 10-18 of the differentiation protocol. Peroxisomes were visualized with the PMP70 antibody, nuclei were stained with Hoechst (10μg/ml), and neuronal differentiation was visualized with β-3 tubulin antibody. Representative images are shown, Scale bar - 5μm. **(D)** Quantitative PCR of the neuronal markers in the 12-day differentiated NPCs. Quantification shows the relative expression of differentiated/non-differentiated markers, expressed as 2^-ΔΔCt^, mean± SEM. **(E)** Confocal microscopy of peroxisomes in the non-differentiated and 12-day differentiated NPCs in control and Go6983 1μM conditions, Go6983 was added on days 1-12 of the differentiation protocol. Peroxisomes were visualized with the PMP70 antibody, nuclei were stained with Hoechst (10μg/ml), and neuronal differentiation was visualized with β-3 tubulin antibody. Representative images are shown, Scale bar - 5μm, inlet - 1μm. **(F)** Confocal microscopy of peroxisomes in the neuronal terminals of the 18-day differentiated SH-SY5Y and 12-day differentiated NPCs in control and Go6983 1μM conditions. Peroxisomes were visualized with the PMP70 antibody, and neuronal terminals were visualized with β-3 tubulin antibody. Representative images are shown, Scale bar - 5μm.

**Figure S8.**
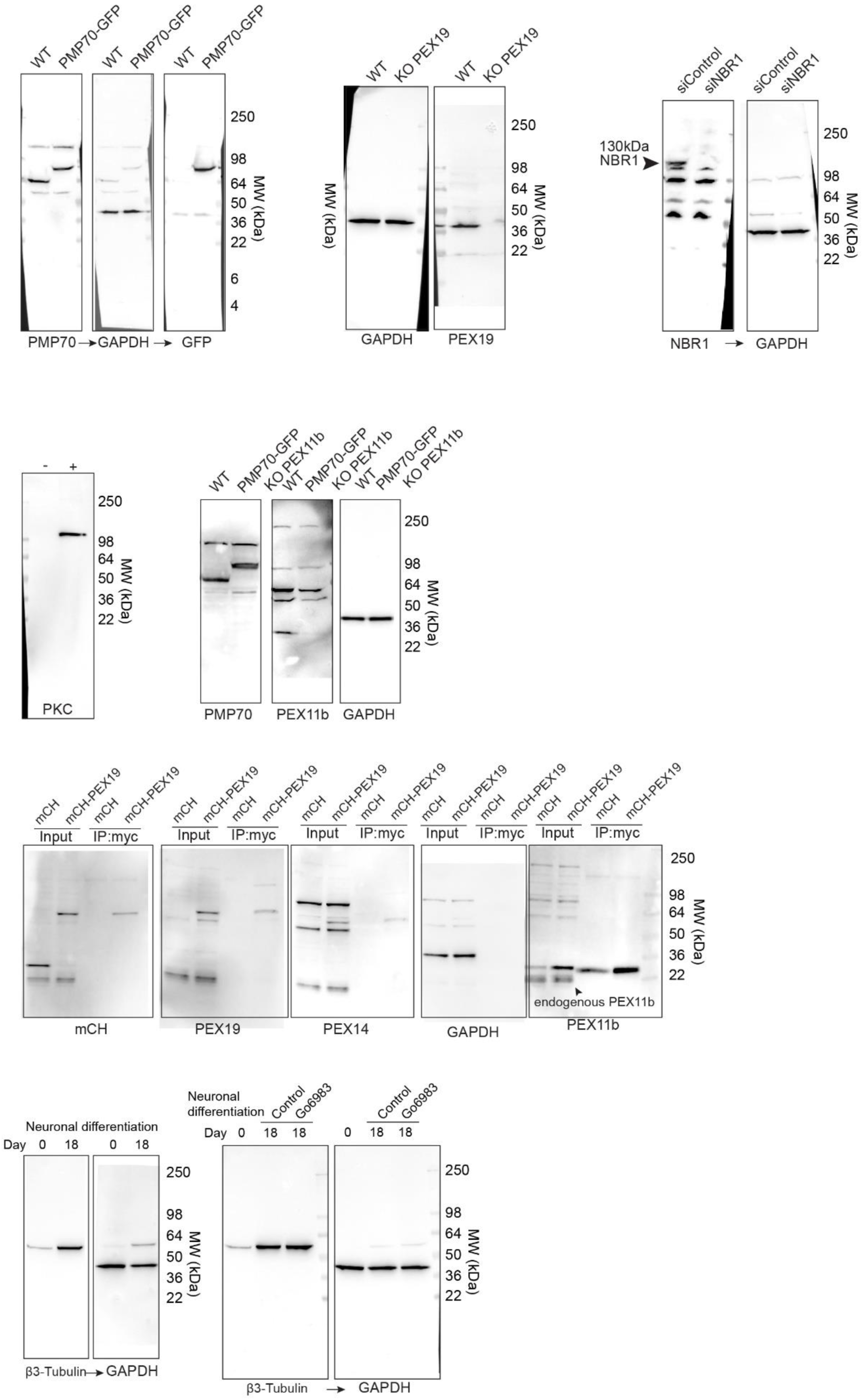
Western blots

**Figure S9.**
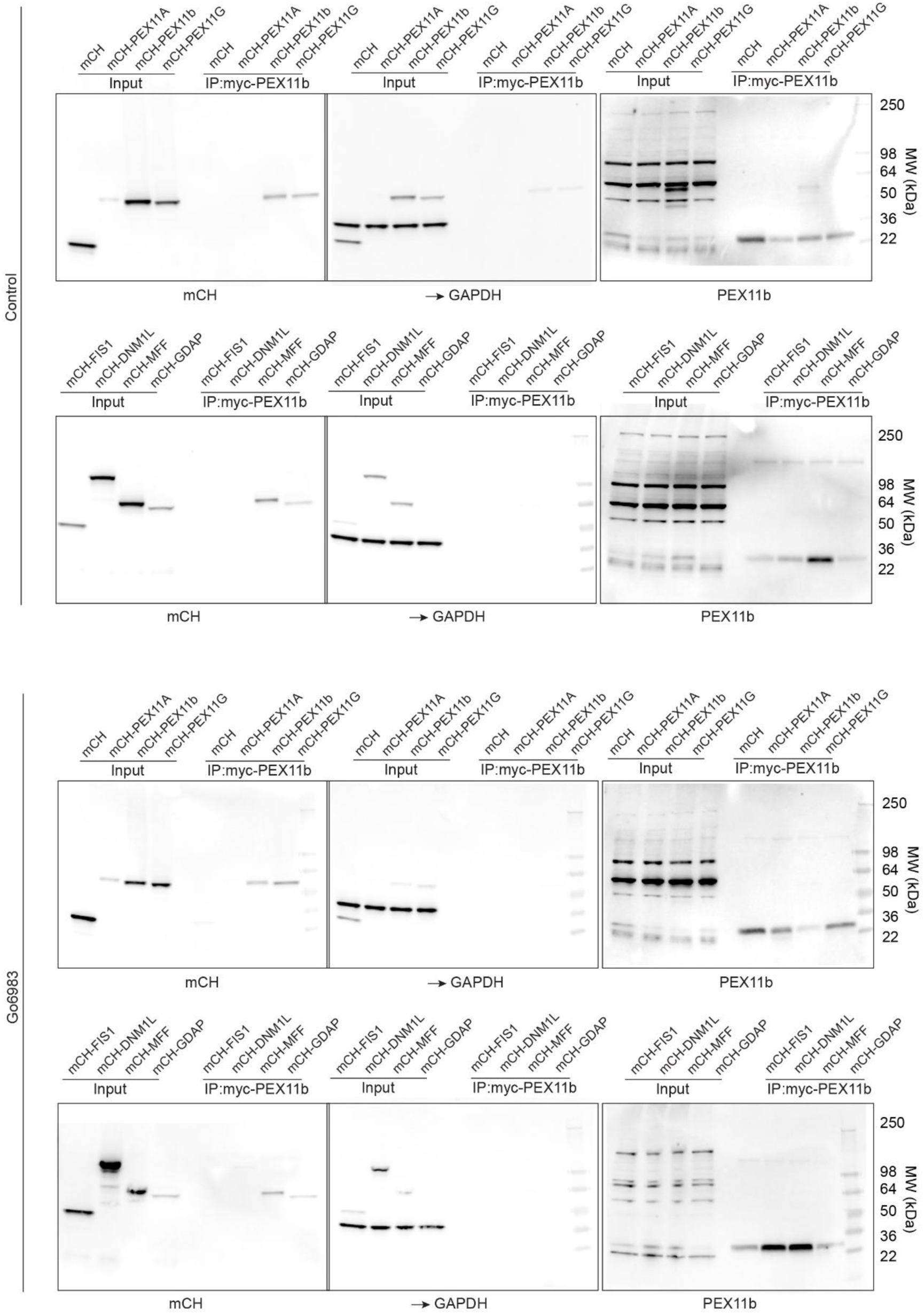
Western blots

**Figure S10.**
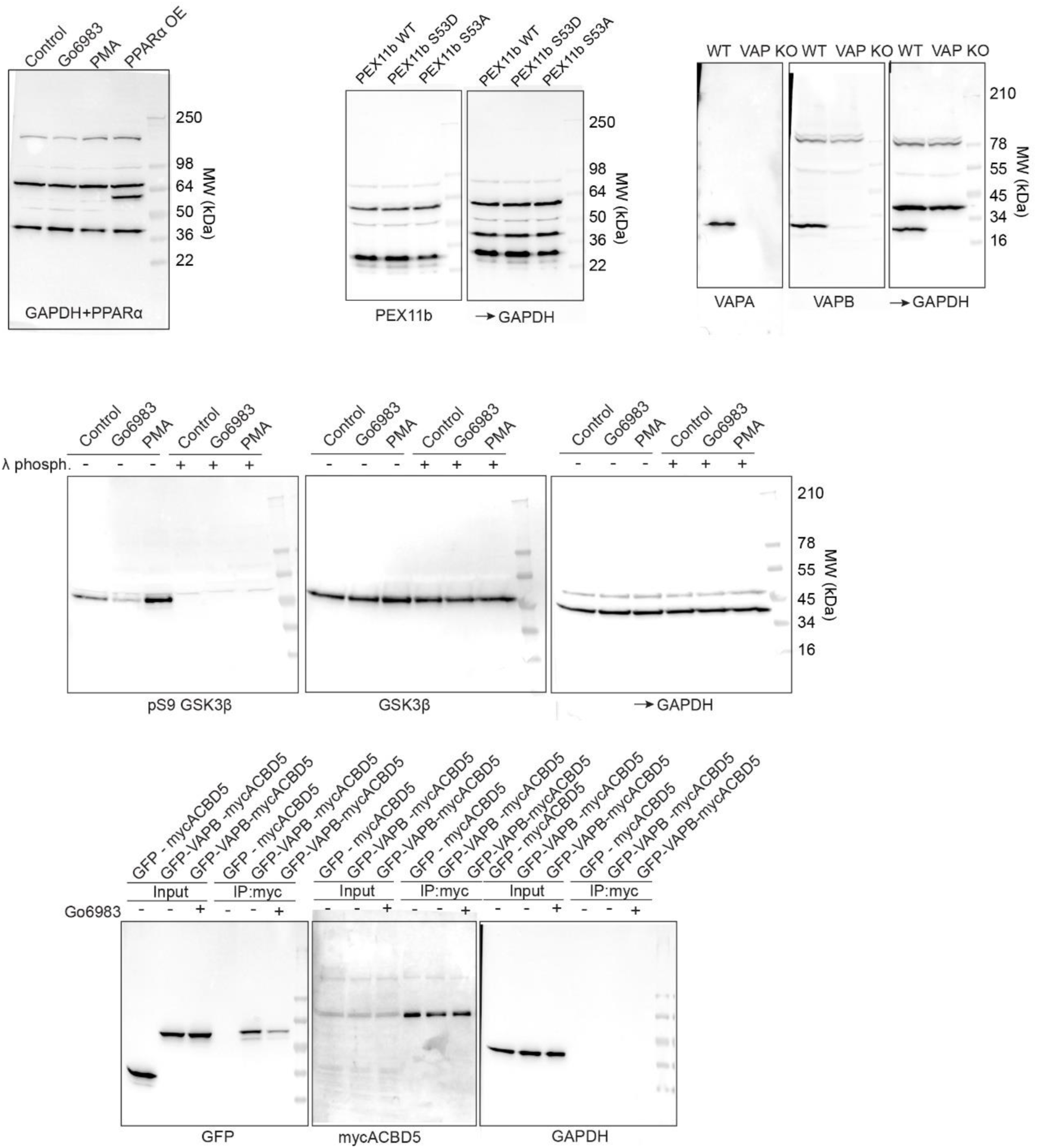
Western blots

